# An Indole Dimer Antifungal Metabolite from a Rice Seed Endophyte Inhibits Ergosterol Biosynthesis in Fungal Pathogens

**DOI:** 10.64898/2026.01.05.697688

**Authors:** Santosh Kumar Jana, Supriya Bhunia, Dipanjan Ghosh, Debashmita Guha, Shrodha Mondal, Himadri Sekhar Sarkar, Samudra Gupta, Subhas Samanta, Prithidipa Sahoo, Sukhendu Mandal

**Author notes:** **To whom correspondence should be addressed:**, **Corresponding Author:** Sukhendu Mandal, **Address:** Laboratory of Molecular Bacteriology, Department of Microbiology, University of Calcutta, 35, Ballygunge Circular Road, Kolkata, 700019, India.

## Abstract

The increasing prevalence of fungal phytopathogens and the widespread emergence of fungicide resistance necessitate alternative antifungal strategies with reduced environmental impact. Here, we report the isolation and characterization of a novel antifungal metabolite, SM06, produced by the rice seed-associated endophytic bacterium *Phytobacter* sp. RSE02. SM06 exhibited broad-spectrum antifungal activity against plant and human pathogenic fungi, including *Curvularia lunata*, *Fusarium oxysporum*, and *Candida albicans*. In vitro assays and micromorphological analyses revealed that SM06, an indole dimer, disrupts fungal cell membrane integrity, while in planta experiments demonstrated significant suppression of brown leaf spot disease in tomato and rice. Molecular docking suggested that SM06 binds to lanosterol 14α-demethylase (ERG11), a key enzyme in fungal sterol biosynthesis. Consistent with this prediction, LC–MS–based analyses confirmed a significant reduction in ergosterol content in SM06-treated fungal cells. Together, these findings identify SM06 as a biologically active antifungal metabolite produced by a plant-associated bacterium and highlight its potential application in sustainable fungal disease management.

**IMPORTANCE:** Fungal diseases cause major losses in crop-production and contribute to the growing challenge of antifungal resistance, underscoring the need for sustainable alternatives to chemical fungicides. This study identifies SM06, a novel indole dimer produced by the rice seed endophyte *Phytobacter* sp. RSE02, with strong antifungal activity against economically important plant pathogens and clinically relevant fungi. Through integrated chemical, cellular, and in planta analyses, we demonstrate that SM06 disrupts fungal membrane integrity by inhibiting ergosterol biosynthesis. The compound is biocompatible, stable, and effective in plant disease suppression, highlighting its translational potential for crop protection. These findings reveal seed endophytes as an important yet underexplored sources of antifungal metabolites and provide a mechanistic foundation for developing eco-friendly biocontrol strategies with implications beyond agriculture.

**Graphical Abstract:** 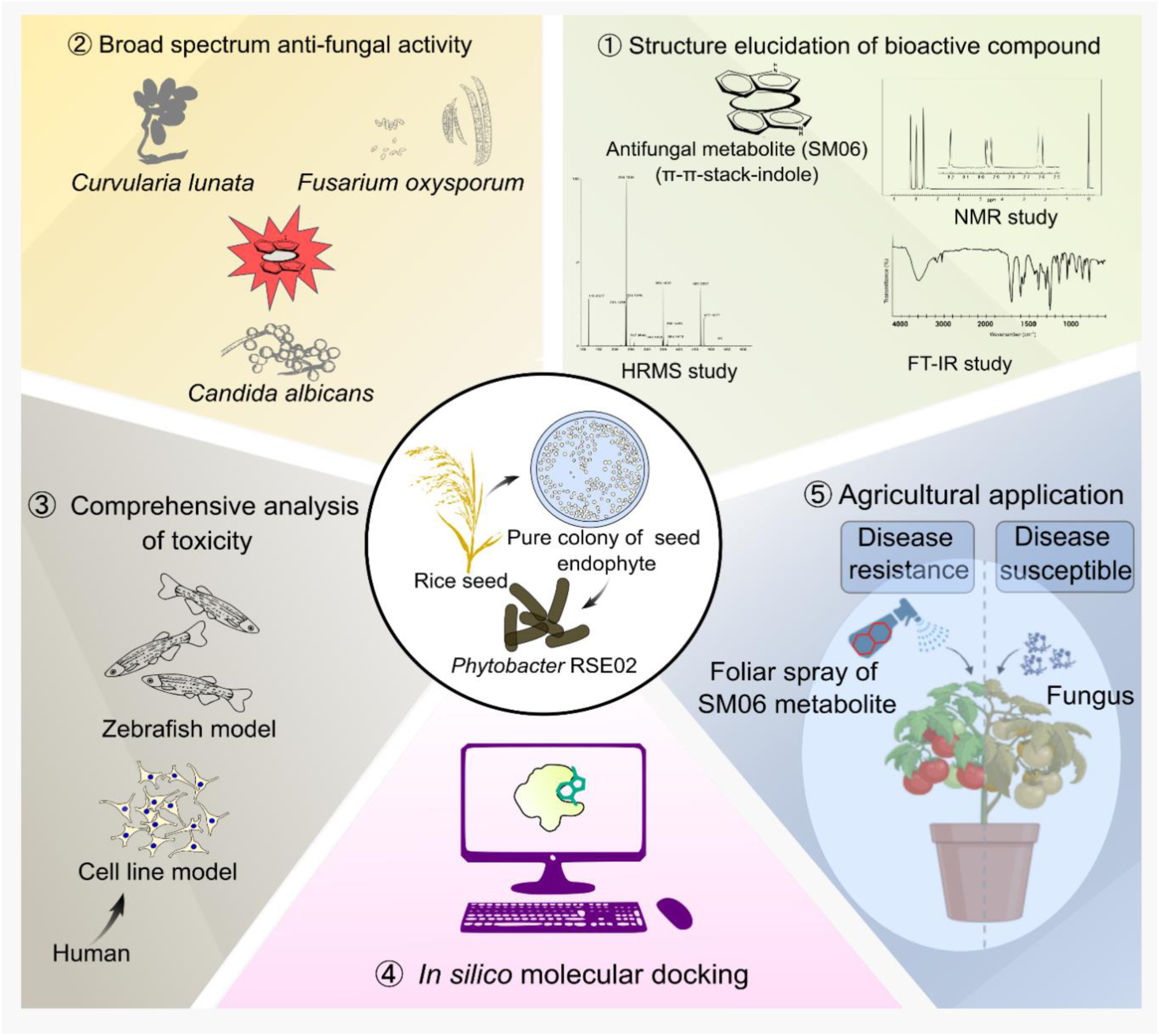

## INTRODUCTION

Ensuring sustainable agricultural productivity has become a global priority in front of rising food demand and increasing environmental pressures associated with conventional farming practices [1]. Among the major constraints to crop production, fungal phytopathogens represent a persistent and widespread threat, contributing to substantial pre- and post-harvest losses, contamination of food commodities with mycotoxins, and reduced market value. It is estimated that fungal diseases alone account for yield losses exceeding 20% across several economically important crops worldwide [2,3]. Synthetic fungicides such as carbendazim and thiophanate-methyl have been extensively employed to manage these infections [4]; however, their long-term use has resulted in significant drawbacks, including environmental persistence, toxicity to non-target organisms, and increasing regulatory restrictions. Moreover, repeated exposure has accelerated the emergence of fungicide-resistant pathogen populations, diminishing their effectiveness over time [5,6]. Resistance has been widely reported in genera such as *Fusarium*, *Botrytis*, *Alternaria*, and *Penicillium*, often arising through point mutations in target enzymes, overexpression of efflux transporters, alterations in target site architecture, or enhanced metabolic detoxification pathways [7].

In response to these limitations, biological control strategies have gained increasing attention as environmentally compatible alternatives for fungal disease management [8]. The plant-associated bacteria are particularly attractive as they synthesize a wide array of secondary metabolites with antifungal properties [9]. Endophytic bacteria, which reside asymptomatically within plant tissues, contribute to host fitness by promoting growth and enhancing tolerance to biotic and abiotic stresses. Members of the family Enterobacteriaceae, including species of the genus *Phytobacter*, have been reported to exhibit plant growth-promoting traits and antagonistic activity against phytopathogens [10]. The application of such beneficial endophytes aligns with sustainable agriculture goals by reducing reliance on synthetic fungicides while supporting crop productivity and soil health [11]. Consequently, exploring underexplored microbial taxa such as *Phytobacter* for antifungal metabolite production represents a promising avenue for the development of next-generation biocontrol agents [12].

A key attribute of endophytic microorganisms is their capacity to produce structurally diverse bioactive secondary metabolites, including alkaloids, polyketides, terpenoids, peptides, phenolics, and lipopeptides, many of which possess strong antimicrobial activities [13,14]. Endophytic fungi have historically yielded several clinically and agriculturally relevant antifungal compounds, such as griseofulvin and trichothecenes. Similarly, bacterial endophytes produce antifungal metabolites including pyrrolnitrin, phenazine-1-carboxylic acid, and various biosurfactants, some of which also elicit induced systemic resistance in plants [15]. These observations highlight endophytes as prolific reservoirs of chemically and functionally diverse antifungal agents. Such agents also have very specific targets present among respective target organisms. The uniqueness of these targets make an antifungal agent specific towards a group without having toxicity against other organisms. At the molecular level, ergosterol biosynthesis remains one of the most effective targets for antifungal intervention. The cytochrome P450 enzyme lanosterol 14α-demethylase (ERG11) catalyzes a critical step in this pathway by removing the 14α-methyl group from lanosterol and is the primary target of azole-class antifungals such as fluconazole and itraconazole [16]. However, resistance to azoles frequently arises through mutations in ERG11 that reduce inhibitor binding efficiency [17]. Heterocyclic scaffolds, particularly indole-based compounds, are of interest for antifungal drug design due to their ability to engage in π–π stacking interactions with the heme group and aromatic residues within the ERG11 active site [18]. In silico docking approaches provide valuable insights into ligand orientation, residue-level interactions, and binding affinities, thereby guiding experimental validation and aiding in the identification of compounds with improved efficacy and resistance resilience. These considerations emphasize the need for environmentally benign antifungal agents that operate through well-defined molecular mechanisms [19].

In this study, we investigated *Phytobacter* sp. RSE02, a rice seed-associated endophytic bacterium previously identified as a plant growth-promoting strain [20], for its antifungal metabolite production. Preliminary assays revealed that culture supernatants of RSE02 exhibited broad-spectrum antifungal activity against multiple fungal pathogens [20]. Here, we report the purification and structural characterization of a di-indole antifungal compound, designated SM06, produced by RSE02. We further examine its antifungal efficacy, effects on fungal cell integrity, and potential mechanism of action, including interaction with ERG11. The identification of SM06 highlights the untapped metabolic potential of seed-derived bacterial endophytes and underscores their relevance as sources of biologically active antifungal metabolites for sustainable disease management.

## MATERIALS AND METHODS

### Preliminary screening for antimicrobial property

In this study, we have investigated the bioactive compounds produced by the endophytic bacterium *Phytobacter* sp. RSE02 (GenBank accession no. OM403299), which was isolated from viable rice seeds, as previously described by Jana et al. 2023 [20]. Using Mueller-Hinton (MH) agar media and the conventional agar-diffusion method, the antimicrobial activity of the strain RSE02 was evaluated [21]. The test organisms used in this method were *S. aureus* MTCC 96, (Gram-positive), *E. coli* MTCC 1687, (Gram-negative), plant fungal pathogen *Curvularia lunata* (MTCC accession no. 8018) and *Fusarium oxysporum* (MTCC accession no. 284), human fungal pathogen *Candida albicans* (MTCC accession no. 183) and *Rhizopus stolonifer* (MTCC accession no. 4886). 2-3 days old RSE02 culture supernatant was used to check the antibiosis against the aforesaid strains. The plates were incubated at 37°C for 3-4 days.

### Optimization of SM06 production

With different media conditions, incubation times, the best way to production of the antifungal molecule produced by RSE02 has been investigated. While growth factors like inoculum (1%) and best production time (16 h), shaking speed (180 rpm), and temperature (30°C) were held constant.

### Production, extraction, and purification of SM06

Overnight grown culture of RSE02 (1%, v/v) was inoculated in 2 L of sterile Luria-Bertani (LB) medium, and incubated at 30°C for 16 h with 180 rpm of shaking. The culture medium was centrifuged at 13,000 rpm for 15 min to collect the supernatant. An equal volume of ethyl acetate was added to the supernatant, and the mixture was shaken properly to ensure thorough extraction of the active compound [22]. The active compound was collected after drying the organic phase using rotary evaporator. After resuspension with 200 μL ethyl acetate, the fraction was run in thin layer chromatography (TLC) plate (silica gel 60 F254) using PET: ethyl acetate (9:1, v/v), Rf = 0.40). To obtain the required product, silica gel column chromatography (60-120 mesh size) was applied to the crude product with the same solvent eluent ratio. The solvent was removed from the eluted fractions using a rotary evaporator, the pure product was obtained as ash-gray solid. The 2 L extract solutions yielded approximately 42 mg of pure product. The product was then further employed for its chemical and functional characterization.

### Antimycotic activity of RSE02 crude extract and SM06 against fungal pathogens

To evaluate the antifungal activity of RSE02 crude extract and the SM06 compound against *C. lunata*, *F. oxysporum*, *C. albicans*, and, *R. stolonifer* we followed the methodology outlined in Ruhil et al. (2013) [23]. Briefly, each fungal suspension (100 μL) was uniformly spread onto Mueller-Hinton agar plates. A volume of 4 μL containing RSE02 crude, SM06, Nystatin, Cycloheximide, or biogenic silver nanoparticles (AgNPs) was individually applied to the inoculated plates. After incubation at 37°C for 3 days, the diameters of inhibition zones were measured.

To assess the concentration-dependent inhibition of *C. lunata* biomass by SM06, sterile 2 mL culture tubes were each filled with 900 µL of sterilized culture medium. Fresh mycelial inocula (100 µL) were added to all tubes. Subsequently, SM06 was added at increasing volumes 5, 10, and 20 µL from a 15 µg/mL stock solution. Cycloheximide and Nystatin were included as positive controls. Following 7 days of incubation at 37°C, the mycelial growth inhibition was quantified by comparing biomass between treated and control samples, expressed as percentage inhibition of mycelial growth [24]. The antibiosis against both Gram-Positive and Gram-negative bacteria were also tested taking *Staphylococcus aureus*, and *Escherichia coli*, respectively as standard strains [20, 21].

### Structural elucidation of SM06

The solvents were distilled and dried following the standard procedures [25]. High-resolution mass spectrometry (HRMS) was carried out using a Waters, XEVO G2-XS QTOF instrument by using acetonitrile (ACN) as the solvent. Bruker Avance 300 (300 MHz) and Bruker Avance Neo HD 400 (400 MHz) were used to record the ^1^H and ^13^C NMR. For NMR spectra, CDCl_3_ was used as solvent and TMS as an internal standard. Chemical shifts are expressed in δ ppm units and ^1^H–^1^H, ^1^H–^13^C coupling constants in Hz. The following abbreviations describe spin multiplicities in ^1^H NMR spectra: s = singlet; d = doublet; t = triplet; m = multiplet. UV spectra were recorded on Ava Spec-ULS2048L-USB2-UARS spectrometer with temperature-controlled cuvette holder. FTIR data were collected using FTIR-7600S (Lambda-Scientific) spectrometer.

### Assessment of antifungal activity

#### Sporicidal and mycelium disruption efficacy of SM06 against *C. lunata*

To evaluate the inhibitory effect of SM06 on spores and mycelia of *C. lunata*, assays were conducted in sterile 2 mL culture tubes, each containing 900 µL of sterilized culture medium [26]. A working concentration of SM06 (15 µg/mL) was prepared. 100 µL of *C. lunata* spore or mycelium suspension was introduced in each tube. Tubes containing only the spores or mycelia suspension and devoid of SM06 served as negative controls. Following inoculation, all tubes were incubated at 37°C for 16 h. After incubation, spore germination and hyphal (mycelial) morphology were assessed microscopically using an Olympus cellSens imaging system and scanning electron microscope (SEM) system developed by Carl Zeiss Microscopy GmbH. We also performed time kill kinetics after certain time intervals (0, 1.5 and 2.5 h), fungal hyphae from treated and control samples were examined under light microscope (Olympus cellSens) to observe morphological alteration [21].

We assessed membrane integrity of the test fungal strain in both compound-treated and untreated hyphae by staining with propidium iodide (10 µg/mL), a membrane-impermeant fluorescent dye, and visualized them using fluorescence microscopy [27]. The fluorescence intensity of individual hyphal segments was then quantified using ImageJ software (version 1.54g).

### Membrane integrity assessment via extracellular protein leakage

By monitoring the leakage of cell constituents, such as proteins, into the cell suspension, the integrity of the cell membrane was assessed [28]. Centrifugation at 6000 rpm for 5 min was used to collect the actively growing fungal mycelial pellet. This pellet was washed with phosphate-buffered saline (PBS) (pH 7.0) for 3 times. To assess membrane integrity, the pellet was treated with the minimum inhibitory concentration (MIC) of SM06 (15 µg/mL) and incubated over multiple time intervals (0, 1, 4, 8, 12, 16, 20, and 24 h). At each time point, 1 mL of the suspension was collected and centrifuged at 6,000 rpm for 10 min. The total proteins present in the supernatant were measured through the Bradford assay procedure [29]. Each treatment and measurement was performed in triplicate to ensure reproducibility and statistical reliability.

### Antifungal efficacy of SM06 in combination with Nystatin

To determine the combinatorial effect of SM06 with frequently used antifungal antibiotics like nystatin, a checkerboard assay method following the standard protocol as mentioned in Sardana et al. (2021) [30]. Antibiotics used in the experiment ranged from MIC to 1/6 MIC equivalent. The final antibiotic concentrations were: Nystatin 0.08, 0.16, 0.32, 0.64, 1.28, 2.56 and 5.12 μg/mL; and SM06 0.8, 1.6, 3.2, 6.4, 12.8, 25 and 50 μg/mL. Experiments were done in a 96-well plate with 100 μL of MH broth in each well. Each well has 40 μL of MH broth, 50 μL of freshly diluted test organism, 5 μL of the first test compound (SM06), and 5 μL of the test antibiotic. 1 × 10⁶ spores/mL was the final inoculum, additionally, each well contains an inoculum of the test pathogen *C. lunata* together with the specific individual antibiotics. The cells were incubated for 48 h at 30°C. The fractional inhibitory concentration index (FICI) was used to quantify in vitro interaction and the formula of FICI calculation is (MIC of drug A in combination/MIC of drug A alone) + (MIC of drug B in combination/MIC of drug B alone). FICIs were interpreted as follows: < 0.5, synergy; 0.5-0.75, partial synergy; 0.76-1.0, additive effect; 1.0-4.0, indifference; and > 4.0, antagonism. The varying levels of synergy between two given compounds were determined [30]. The experiment was performed in triplicate.

### In vitro toxicity testing and localization profiling of SM06 compound in zebrafish embryos and mammalian cells

#### Zebrafish maintenance

Adult zebrafish of the AB strain with ages between 3-6 months were used in this study for spawning. Fish were maintained in customised aquarium tanks at a density of about 5 fish per litre at 28℃ water temperature and fed three times daily. Collection of zebrafish embryos was performed after natural mating and maintained in E3 medium at a temperature of 28°C up to 4-day post-fertilisation (dpf). All zebrafish husbandry and experiments were conducted in accordance with institutional and national animal ethics guidelines and were approved by the Committee for the Purpose of Control and Supervision of Experiments on Animals (CPCSEA). (Approval No. IAEC-04/BIOCHEM/CU/02-2022/03-SH)

### Generation of germ-free zebrafish embryos and treatment with SM06

The generation of germ-free embryos and evaluation of the effect of SM06 on zebrafish embryos were done as previously described with minor modifications [31]. Briefly, after 24 hpf (hours post-fertilization), the embryos were dechorionated and divided into two groups: Conventionally raised (CV) and germ-free (GF). The CV larvae were maintained at room temperature while the GF embryos underwent sterilization by immersion in sterilized Gentamicin (100 µg/mL) for 1 h, followed by treatment in 0.003% hypochlorite solution and subsequently washed in sterile E3 medium under laminar hood to ensure sterility. The non-lethal dose was determined by exposing 24 hpf zebrafish embryos to three metabolite concentrations according to the *in vitro* dose (50 µg/mL, 100 µg/mL, and 250 µg/mL).

A total of ten zebrafish embryos at 24 hpf were placed in a petri dish containing 3 mL of E3 medium supplemented with SM06 (50 µg/mL, 100 µg/mL, and 250 µg/mL) and exposed for 3 days with the medium renewed daily. Embryos exposed to 0.1% DMSO in E3 medium were served as control group.

### Fluorescence microscopic analysis of embryonic gut lumen

The 96 hpf zebrafish embryo after exposure were anesthetized with tricaine and mounted on the grooved slide using 0.1% Methyl cellulose (M7027, Sigma, St. Louis, MO, USA) as described by Mukherjee et al. (2025) [32] and photographed by using an OLYMPUS BX53F2 fluorescent microscoped (Olympus, Tokyo, Japan) under 335 nm wavelength filter to observe the fluorescent intensity of the bacteria metabolite in the gut lumen of the embryo.

### Mammalian cells maintenance

HeLa cell lines were prepared from a continuous culture in Dulbecco’s Modified Eagle’s Medium (DMEM, Sigma Chemical Co., St. Louis, MO) supplemented with 10% fetal bovine serum (Invitrogen), penicillin (100 μg/mL), and streptomycin (100 μg/mL). After the cells reached the logarithmic phase, the cell density was adjusted to 1.0 × 10^5^ per culture dish in culture media and then used to inoculate with 1.0 mL (1.0 × 10^4^ cells) of cell suspension in each glass bottom dish.

We have performed MTT assay to evaluate the cytotoxicity of compound SM06 on the HeLa cell line by a standard protocol [33]. A 96-well polystyrene-coated plate with 100 μL of 7500 cells were incubated for 24 h at 37°C in an incubator having 5% CO_2_ and 95% air. Each SM06 dilution was mixed (1x, 2x, 3x and 4x MIC μg/mL) (MIC is 15 μg/mL) in the medium taken in wells of a 96-well plate. Each dilution point was taken in triplicate. 20 μL MTT reagent was added in each well and incubated for an additional 3 h. After formation of the purple formazan crystals, 150 μL of MTT solvent was added in each well of the microplate and scanned using a microplate reader Synergy H1 (Agilent-Biotek) at 590 nm. The reading from the wells which contained medium, SM06, and MTT reagent but no cells were considered as blank.

### *In-silico* interactions between indole dimer and ERG11 complex

The *in-silico* molecular docking study was performed between indole dimer and ERG11 complex with the CB Dock tool [34]. CB Dock is an online docking tool, while the docking algorithm is programmed by Auto Dock Vina [34]. The crystal structure of the ERG11 protein was retrieved from the Protein Data Bank (PDB: 5EQB, https://www.rcsb.org/structure/5EQB). However, the established structure of the ligand molecules [such as normal indole, fluconazole (positive control), and ergosterol (substrate of ERG11)] was retrieved from the PubChem database (https://pubchem.ncbi.nlm.nih.gov/). The 3D structure of the stacked indole dimer was designed using Avogadro software [35]. The energy-minimized structure of the ligand molecules was used for *in silico* docking analysis. The energy minimization of the ligands was performed with Avogadro software using the MMFF94 force field [36]. The docking result was analyzed using PyMOL (Schrodinger, v 2.5.8) [37] and Discovery Studio Client software (v 24.1.0.23298).

### *In-silico* ADMET prediction of the indole and indole dimer

The ADMET and physicochemical properties of the indole and indole dimer were collected from the ADMET-AI prediction tool server (https://admet.ai.greenstonebio.com/). It is a web-based platform to find out the ADMET property of the unknown compound [38].

### Estimation of ergosterol from *C. lunata* upon SM06 treatment

To evaluate the time-dependent effects of SM06 on ergosterol biosynthesis in *C. lunata* mycelia, ergosterol was extracted using a chloroform:methanol solvent system following the protocol described by Wilkes, 2023 [39]. The resulting lipid extracts were analyzed by LC–MS to quantify ergosterol abundance across treatment time points using a Waters Xevo-XS Q-TOF (ESI)–UPLC system, with separation on a reverse-phase C18 column and identification in positive ion mode based on characteristic m/z values from extracted ion chromatograms.

### In planta antagonistic activity of SM06 against *Curvularia* in tomato and rice plant

#### Detection of indole dimer synthesis by RSE02 inside tomato plants

We investigated whether *Phytobacter* sp. RSE02, when present inside tomato plants can synthesize SM02. We first monitored the localization of the RSE02 (labelled with red-fluorescent protein) within tomato plants following the laboratory standard described in Jana et al. (2025) [40]. We further aimed to perform the LC-MS analysis to detect SM06 from tissues of healthy plant and *C. lunata* infected plant with RSE02 treatment. Harvested tissues were extracted with ethyl acetate. Following extraction, a rotary evaporator was used to collect and dry the active organic phase and then run through LC-QTOF-MS (Agilent), with SM06 spiked in set as our standard control for indole dimer detection. Retention time, mass/charge (m/z) values, and peak shapes for SM06 were established under these conditions [41].

### Pathogenic challenge of tomato and rice with *Curvularia* species

*C. lunata* was cultivated on potato dextrose broth (PDB) for 10 days at 30℃ under darkness to yield spores. Thirty-day old tomato and rice plants were infected with the pathogen using the punch inoculation method, as described by Wang et al. (2018) [42]. Spores were used at a concentration of 5×10^5^ /mL for punch inoculation. After lightly pounding each plant leaf with a puncher, 5 µL of spore suspension was added. After that, the spore suspension was kept in place by covering both sides with opaque adhesive tape. The inoculated leaves were photographed twelve days following the inoculation.

### Pre-infection and post-infection treatments with SM06 in *C. lunata* infected tomato and rice plant

After the successful infection of the tomato and rice plants with *C. lunata*, SM06 was applied. SM06 (60 µg/mL) was applied as foliar sprays or root drenches on both plants, focusing on the leaf surfaces where the pathogen tends to thrive. [43, 44]. There was a total of four experimental sets: set-I consisted of plants without pathogen infection; set-II was infected plants without SM06 treatment (positive control); set-III consisted of infected plant with SM06 treatment and set-IV was SM06 pre-treated (from 10 days after seedling germination) plant before pathogen infection. Depending on the severity of the infection, we applied the SM06 compound solutions at regular intervals, preferably during the early stages of pathogen infestation.

### Microscopic evaluation of antibiosis by SM06

For the microscopic visualization of the in-planta antifungal activity of the SM06 against the fungal pathogen, transverse sections of 30-day-old growing roots of rice plants and leaves of tomato plants were prepared and placed on grease-free slides. The section was stained with lactophenol-cotton blue and observed under microscope. Images were captured with OLYMPUS cellSens standard software under 10X magnification [45].

### Measurement of necrotic lesions

H_2_O_2_ accumulation from the necrotic tissues of the tomato leaf-lesions were detected using 3,3’-diaminobenzidine (DAB) staining following Roschzttardtz et al. (2009) [46]. Leaf samples were subsequently washed with water and were submerged in DAB (100 mg DAB, 25 µL Tween 20, 2.5 mL of 200 mM NaHPO_4_ for 100 mL solution) solution. Then the samples were vacuum-infiltrated for 15 min and incubated in the dark for 8 h. The leaves were kept in a water bath at 70℃ for 10 min, and bleaching solution (ethanol: acetic acid: glycerol) was added until the chlorophyll was removed and leaves turned pale or transparent. The samples were washed with water and then imaged.

### Investigation of genetic clusters present in RSE02 genome

The gene clusters associated in the synthesis of secondary metabolite of RSE02 were analyzed using Proksee (https://proksee.ca) and the antibiotics and Secondary Metabolite Analysis Shell (antiSMASH) version 8.0.1, a comprehensive bioinformatics tool designed for the identification and annotation of biosynthetic gene clusters (BGCs) in bacterial genomes [47]. The analysis was conducted with relaxed and loose strictness settings to allow for broader detection of putative biosynthetic gene clusters, including borderline or less-conserved clusters that might be missed under stricter parameters. This approach maximizes sensitivity and captures a more comprehensive secondary metabolite profile.

To enhance cluster identification and comparative analysis, the following similarity search modules were activated, KnownClusterBlast was used to compare predicted gene clusters to experimentally characterized clusters in the MIBiG database to identify known metabolites with similar biosynthetic origins. Whereas, ClusterBlast performs genome-wide comparisons of predicted clusters against a large database of microbial genomes to find related gene clusters in other organisms. SubClusterBlast helps to identify conserved sub-clusters or gene modules shared between biosynthetic clusters, providing insights into cluster modularity and evolution.

### Statistical interpretation

All experiments were performed with a minimum of three replicates, and the results were expressed as the mean ± standard deviation. For the statistical analysis, one-way and two-way analysis of variance (ANOVA), followed by Dunnett’s multiple comparisons test and mean values in each column do not differ significantly at p<0.05. Survival curves were analysed using the Kaplan-Meier method. All experiments were done in triplicate for a total of 10 embryos per condition. All graphs and data were subjected to an analysis of variance test by using the GraphPad Prism 8.0.1 version software and OriginPro 8.0 version software [20].

## RESULTS

### SM06 produced by RSE02 effectively inhibits plant and human pathogenic fungi

SM06 displayed potent antifungal activity against *C. lunata*, *F. oxysporum*, and *C. albicans*, as evidenced by pronounced inhibition zones around wells treated with RSE02 culture suparnatant, purified SM06, and reference antifungals (Fig. 1AI–IV). However, it failed to inhibit the growth of *R. stolonifer*. SM06 also remained non-effective against tested Gram-positive and Gram-negative bacteria. Spore germination assays further revealed extensive germ tube elongation in the untreated controls (Fig. 1BI), whereas SM06 treatment markedly reduced both the spore number and germ tube length (Fig. 1BII). Biomass quantification of *C. lunata* revealed a clear dose-dependent suppression in growth by SM06 (Fig. 1C), comparable to the inhibitory effects of Nystatin and Cycloheximide. Consistent with these findings, disc-diffusion assays on solid plate also confirmed the substantial antifungal potency of purified SM06 in comparison to the DMSO controls (Fig. 1D). Furthermore, SM06 retained full functional activity across 30–90°C, demonstrating high thermal stability (Fig. 1E).

**Fig. 1.**
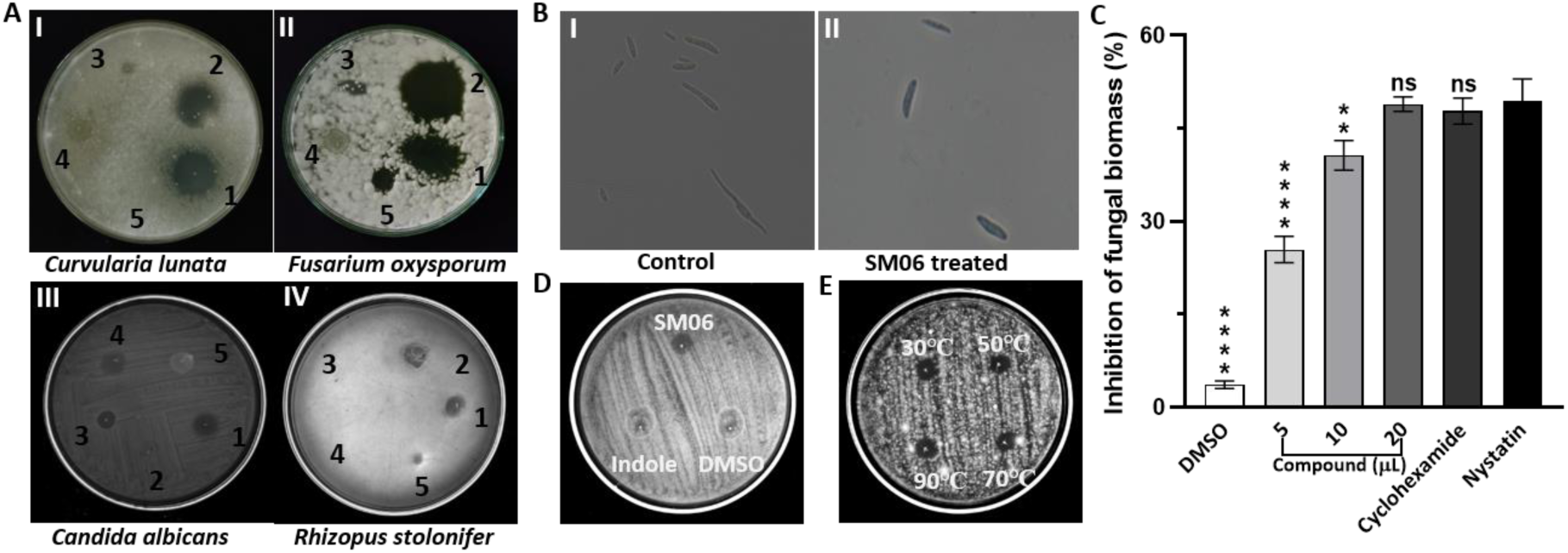
Broad-spectrum antifungal performance and thermal resilience of SM06. A, Antifungal efficacy of SM06 against. I: *Curvularia lunata*; II: *Fusarium oxysporum*; III: *Candida albicans*; IV: *Rhizopus stolonifer* (where 1: Cycloheximide, 2: Nystatin, 3: SM06, 4: Crude SM06 and 5: AgNPs). B, Sporicidal effect of SM06 against *C. lunata* spores. C, Concentration dependent biomass inhibition of *C. lunata* by SM06. D, Comparison study of indole dimer with known indole molecule against *C. lunata*. E, Temperature stability of SM06 compound. Mean values in each column do not differ significantly at p<0.05. Comparison between control and treatment mean values was made by Dunnett’s multiple comparisons test. Where, “****” = p<0.0001, “**” = p<0.01 and “ns” = data is not significant. Data were subjected to analysis of variance (ANOVA one way) test by using the GraphPad Prism 8.0.1 version software.

### SM06 is a novel biomolecule consisting of π–π–stacked indole units

SM06 was obtained as a water-insoluble ash-grey solid and purified by column chromatography using EtOAc:PET (1:9). HRMS analysis showed a dominant [M]^+^ ion at m/z 235.1235 with a secondary peak at m/z 236.1246 (Fig. 2A). The combined ^1^H and ^13^C NMR data (Table 1) established the molecular formula as C₁₆H₁₄N₂. UV–visible and fluorescence measurements revealed absorption and emission maxima at 280 and 335 nm, respectively (Table 1; Fig. S1A–D), which is consistent with an indole chromophore. The IR bands at 1483 and 1597 cm⁻¹ correspond to aromatic C=C stretching, while shifts in the N–H stretching region near 3400 cm⁻¹ indicate hydrogen bonding associated with π–π–stacked indole interactions. A broad absorption at 1974 cm⁻¹ further reflects strong hydrogen bonding between paired N–H groups in the stacked structure (Fig. 2B). Detailed NMR analyses, including COSY, HMBC, DEPT-135, and 1H/13C assignments (Fig. S2–S6), supported an indole-1H–derived scaffold (Fig. 2C). The mass spectrometry data confirmed the presence of an indole dimer (molecular weight 235.1235). Several possible spatial configurations of the dimer were proposed (Fig. S7) and evaluated using DFT calculations (6-31G**; CPCM-H_2_O), which identified pattern 4 as the most energetically favourable structure (Fig. S8).

**Fig. 2.**
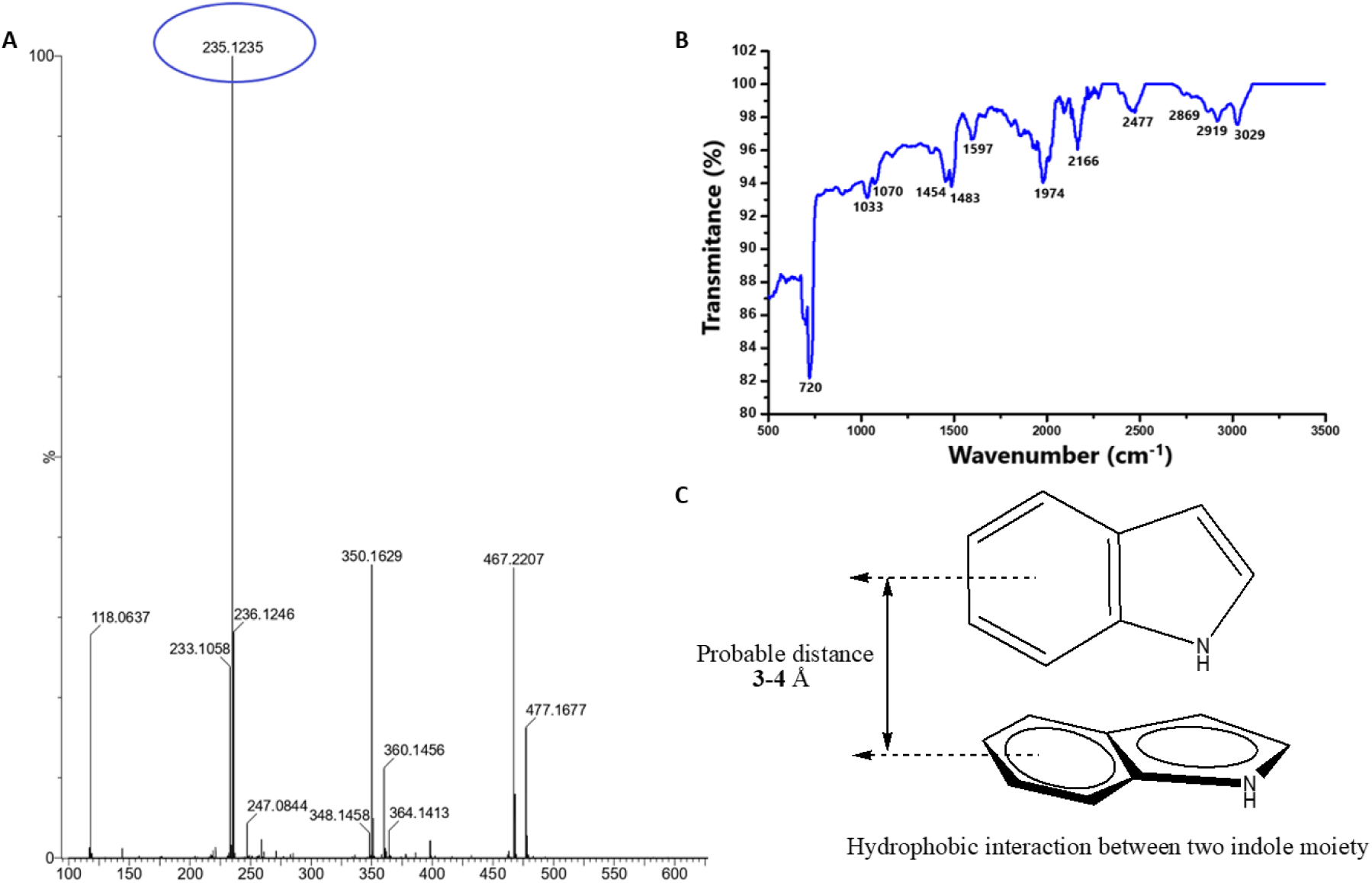
Characterization of compound SM06. A HRMS-spectra of SM06. B FTIR-spectra of SM06 compound. C Structural elucidation of compound SM06.

**Table 1.** Characteristics of the compound SM06.

| Characters | Details |
| --- | --- |
| Appearance | Ash gray solid |
| Molecular formula | C<sub>16</sub>H<sub>14</sub>N<sub>2</sub> |
| Molecular weight | 235.1235 |
| HRMS m/z | (M+H)<sup>+</sup> |
| Found | 235.1235 & 236.1246 |
| Absorbance | 280 nm |
| Emission | 335 nm |
| IR (KBr) cm<sup>-1</sup> | 1483, 1597, 1974, 2166, & 3029 |
| <sup>1</sup>H NMR (400 MHz, CDCl<sub>3</sub>) | δ<sub>ppm</sub> 8.2 (s, 1H), 7.68 (d, 1H, J = 8.0 Hz), 7.41 (d, 1H, J = 8.0Hz), 7.22 (d, 1H, J = 16Hz), 7.14 (t, 2H, J = 12Hz), 6.59 (broad singlet, 1H). |
| <sup>13</sup>C NMR (100 MHz, CDCl<sub>3</sub>) | δ<sub>ppm</sub> 135.8, 127.87, 124.16, 122.01, 120.76, 119.84, 111.05, 102.64 |
| Chemical Name Identified | 1H-indole |

### SM06 leads to the damage of cellular compartments and causes cellular leakage

Scanning electron microscopic (SEM) analysis revealed marked morphological alterations in SM06-treated fungi. Untreated hyphae appeared smooth and structurally intact (Fig. 3AI), whereas SM06 treatment caused swelling, septal distortion, and loss of cell-wall integrity (Fig. 3AII, Fig. S9). Similarly, control spores retained normal surface morphology (Fig. 3AIII), while treated spores showed pronounced surface disruption and fragmentation (Fig. 3AIV). Propidium iodide (PI) staining further confirmed membrane damage where the untreated hyphae exhibited minimal fluorescence (Fig. 3BI–II) as a result of very limited or no entry of PI, but the SM06-treated hyphal showed strong PI uptake at damaged regions (Fig. 3BIII–IV), with significantly elevated fluorescence intensity (Fig. 3C). Protein leakage assays demonstrated a time-dependent increase in extracellular protein over 24 h following treatment at the MIC (15 µg/mL) level of SM06, consistent with progressive membrane permeabilization (Fig. 3D). These results suggest that SM06 disrupts membrane permeability, leading to cytoplasmic leakage and cellular lysis. Antifungal susceptibility testing showed an MIC of 15 µg/mL for SM06 and 1.28 µg/mL for nystatin against *C. lunata*. Combination assays yielded MICs of 1.6 µg/mL (SM06+Nystatin) and 0.64 µg/mL (Nystatin+SM06), corresponding to a fractional inhibitory concentration index (FICI) of 0.60 (Table 2; Fig. S10AI–III), indicating partial synergistic interaction.

**Fig. 3.**
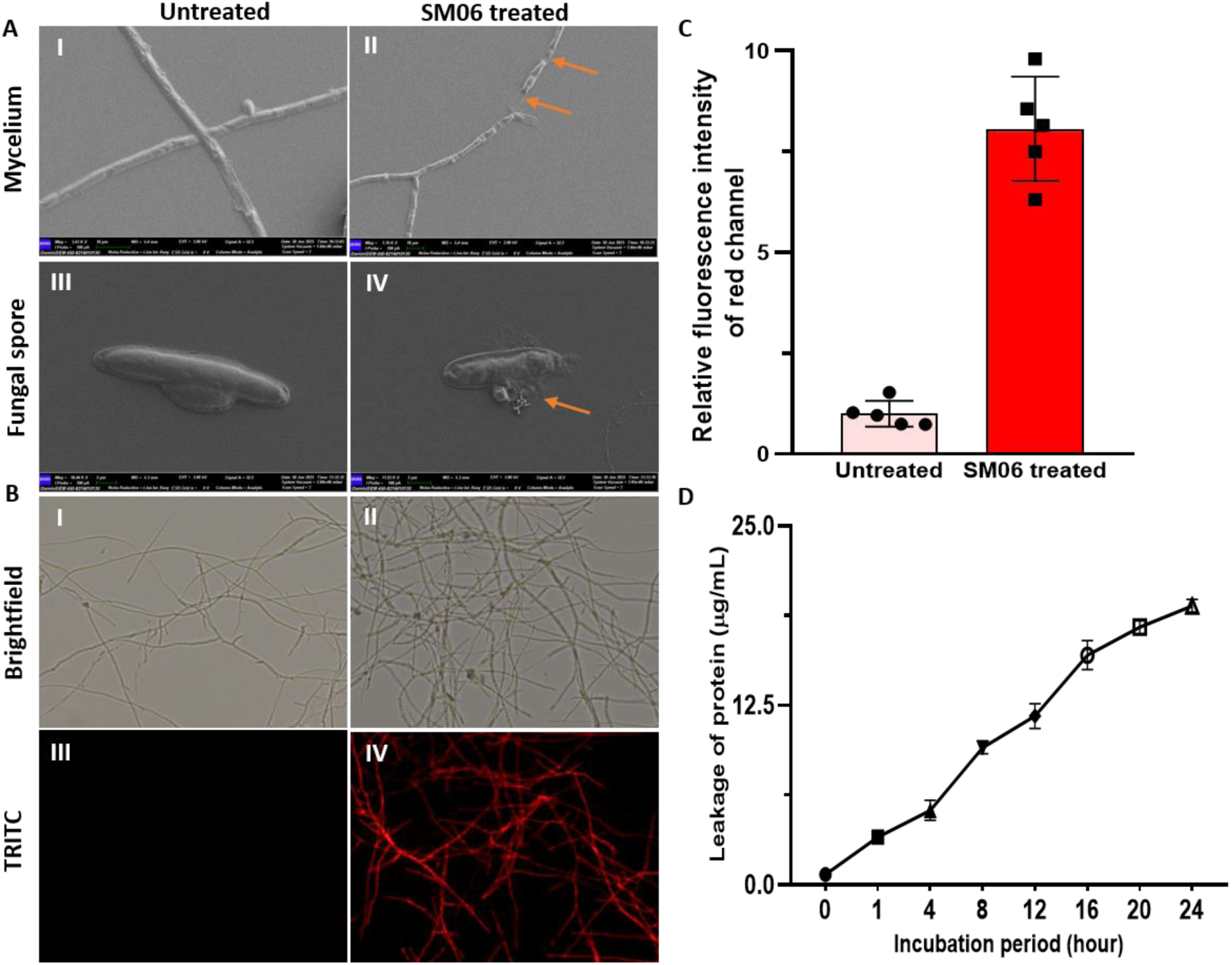
Sporicidal and Mycelial disruption by SM06. AI-IV: Untreated and SM06-treated (15 µg/mL) (for 16 h) *C. lunata* mycelium and spore under SEM. BI-IV: Untreated and SM06-treated (15 µg/mL) *C. lunata* mycelium treated (for 30 min) with propidium iodide (PI). C Relative fluorescence intensity of SM06 treated fungal cell membrane stained with PI. Images from five distinct microscopic fields were analyzed using ImageJ v1.54g software to quantify relative fluorescence intensity. Statistical analysis was performed using GraphPad Prism (version 8.0). Data were expressed as mean ± standard error of the mean (SEM). D: Total protein leakage of *C. lunata* strain treated with SM06 for different time.

**Table 2.**
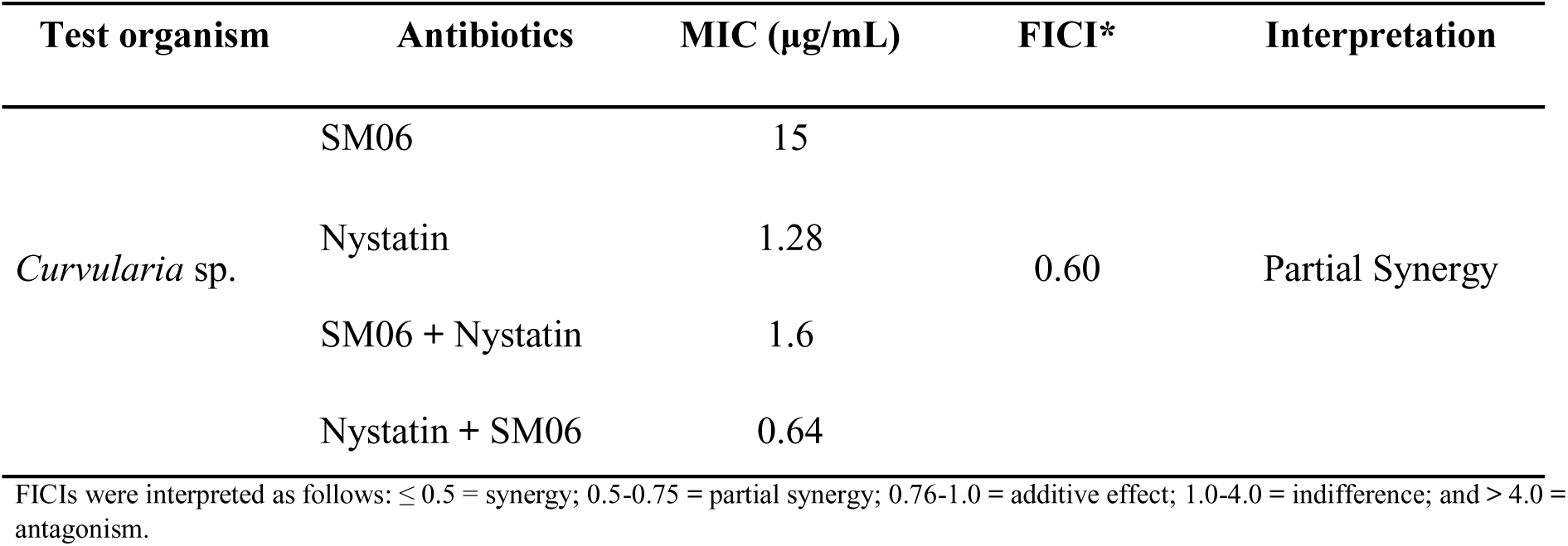
Synergic effect of compound SM06 along with Nystatin against *C. lunata*.

### SM06 is biocompatible and nontoxic for zebrafish embryos and mammalian cell lines

SM06 exhibited no detectable toxicity. Zebrafish embryos exposed to concentrations up to 250 µg/mL from 24–96 hpf showed no significant mortality, and fluorescence imaging indicated the preferential localization of the compound within the intestinal lumen (Fig. 4A–C). Consistently, HeLa cells treated with SM06 at 4× MIC (15 µg/mL) displayed no measurable cytotoxic effects (Fig. 4D). These findings demonstrate that SM06 is biocompatible under the tested conditions and does not elicit adverse effects in zebrafish embryos or mammalian cell lines.

**Fig. 4.**
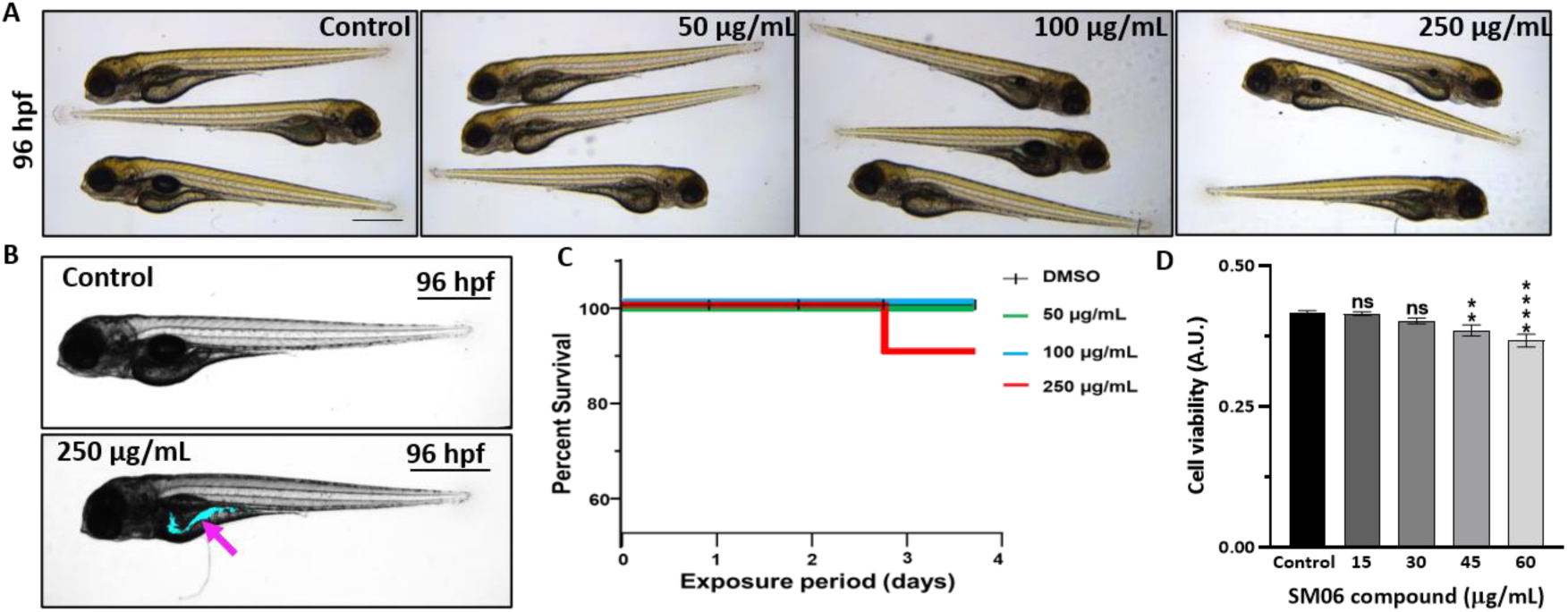
Toxicity and localization profiling of SM06 in zebrafish embryos and mammalian cells. A: for 24 hpf Zebrafish embryos (N=10) were grown in germ-free condition and treated with SM06 (50, 100 and 250 µg/mL) followed by observation after 72 h of treatment at 96 hpf stage. B: Representative image of zebrafish embryo at 96 hpf stage showing localization of SM06 after treating with either DMSO as control (upper panel) or SM06 (lower panel) for 72 h. Visualization of SM06 localized in the embryonic premature gut (marked by arrow) observed at 335 nm excitation under fluorescent microscope (magnification 4x; scale bar 100 μm). C: Kaplan-Meier survival curve of germ-free zebrafish embryos (N=10) after exposure to SM06 (control, 50 µg/mL, 100 µg/mL and 250 µg/mL; each treatment set contains equal amount of DMSO as solvent). D: Effect of SM06 on HeLa cells.

### SM06 binds with fungal ERG11 (Lanosterol 14-α-Demethylase) and inhibits ergosterol biosynthesis

The active site of ERG11 contains a heme cofactor targeted by azole antifungals (e.g., fluconazole) and is positioned within the core catalytic cavity (Fig. 5A–C). Docking analysis showed that lanosterol (substrate for the enzyme ERG11), fluconazole (inhibitor of the ERG11), and indole dimer (SM06) occupied the same binding pocket of ERG11, with a cavity volume of 5267 Å³ (Fig. 5D–H). SM06 exhibited a docking score of –10.5 kcal/mol, which was comparable to that of lanosterol (–10.3 kcal/mol) and stronger than that of fluconazole (–7.0 kcal/mol). Lanosterol formed 20 interactions at the active site (Fig. 5D), whereas fluconazole engaged in 11 interactions (Fig. 5E), which is consistent with their reported binding modes (Fig. 5F; Table S1). SM06 established four key interactions: three hydrophobic contacts with Leu95, Phe241 and Phe384, and a hydrogen bond with Ala125 (Fig. 5G; Table S1). Although the amino acids involved did not fully overlap with the lanosterol interactome, SM06 was bound in the same catalytic pocket as lanosterol and fluconazole (Fig. 5H–I). ERG11 contains a distinct tunnel that ends at its catalytic site and possibly functions as an entry point for lanosterol. The indole dimer SM06 binds to the residues present in the tunnel, impairing the entrance of lanosterol into the catalytic site. Additional cavity morphology comparisons between SM06 and indole are shown in Fig. S11.

**Fig. 5.**
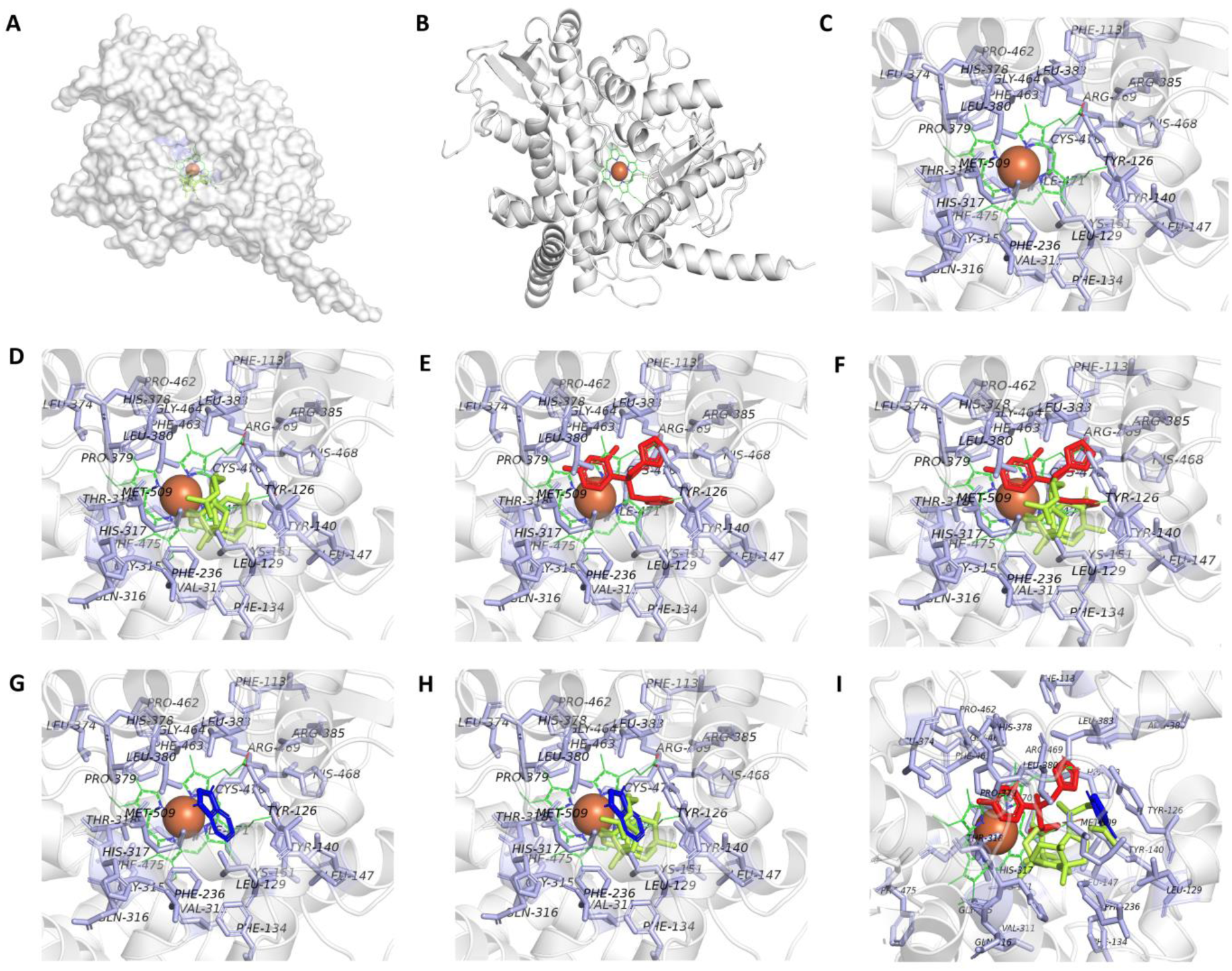
Interaction between indole dimer and ERG11. The docking results were obtained using PyMOL (Schrodinger, v 2.5.8). A and B are the crystal structure of EGR11 (PDB: 5EQB); where, A is surface view of the space-field model and B ribbon view. Both showing the active site (heme group) of the protein along with the iron atom as brick-red color and the porphyrin group in green. C is the zoom-in active site, while D, E, F, G, H and I are ERG11 docked with lanosterol (green color); fluconazole (red color); lanosterol (green color) and fluconazole (red color); indole dimer (blue color); lanosterol (green color) and indole dimer (blue color); and lanosterol (green color), fluconazole (red color) and indole dimer (blue color), respectively. The binding residues were highlighted in light blue.

### Pharmacokinetic and ADMET profiling reveal enhanced oral bioavailability and stability of SM06

In silico physicochemical and ADMET analyses demonstrated the favorable pharmacokinetic properties of SM06 (Table S2). Compared to indole, the indole dimer structure showed increased lipophilicity (∼1.34-fold) and higher predicted plasma protein binding (∼1.61-fold), along with reduced aqueous solubility (20.86% vs. 64.33%, respectively). SM06 also exhibited an elevated TPSA (16.05% vs. 7.95%), indicating greater hydrophilicity. Notably, oral bioavailability was improved (57.58% vs. 52.19%), and the predicted excretion half-life was longer (53.59% vs. 48.89%), suggesting an enhanced metabolic stability. Collectively, these pharmacokinetic features support the superior drug-likeness and biological efficiency of SM06 relative to the parent indole molecule.

### SM06 treatment leads to a significant reduction in ergosterol levels

Ergosterol extraction and LC–MS analysis revealed that treatment with SM06 led to a marked, time-dependent reduction in ergosterol in *C. lunata* (Fig. 6A-C). As shown in the extracted ion chromatograms (m/z 413.2588), the untreated culture displayed a strong ergosterol peak at day 3, whereas equal biomass of SM06-treated samples showed a substantial decrease in peak intensity after 1.5 days and an almost complete depletion after 3 days of incubation (Fig. 6B). Quantitative analysis further corroborated this trend: ergosterol levels were significantly reduced at 1.5 days compared to the untreated control, and by 3 days, ergosterol was nearly undetectable (Fig. 6C). Collectively, these results demonstrate that SM06 effectively inhibits ergosterol biosynthesis in a time-dependent manner, indicating strong interference with the sterol production pathway.

**Fig. 6.**
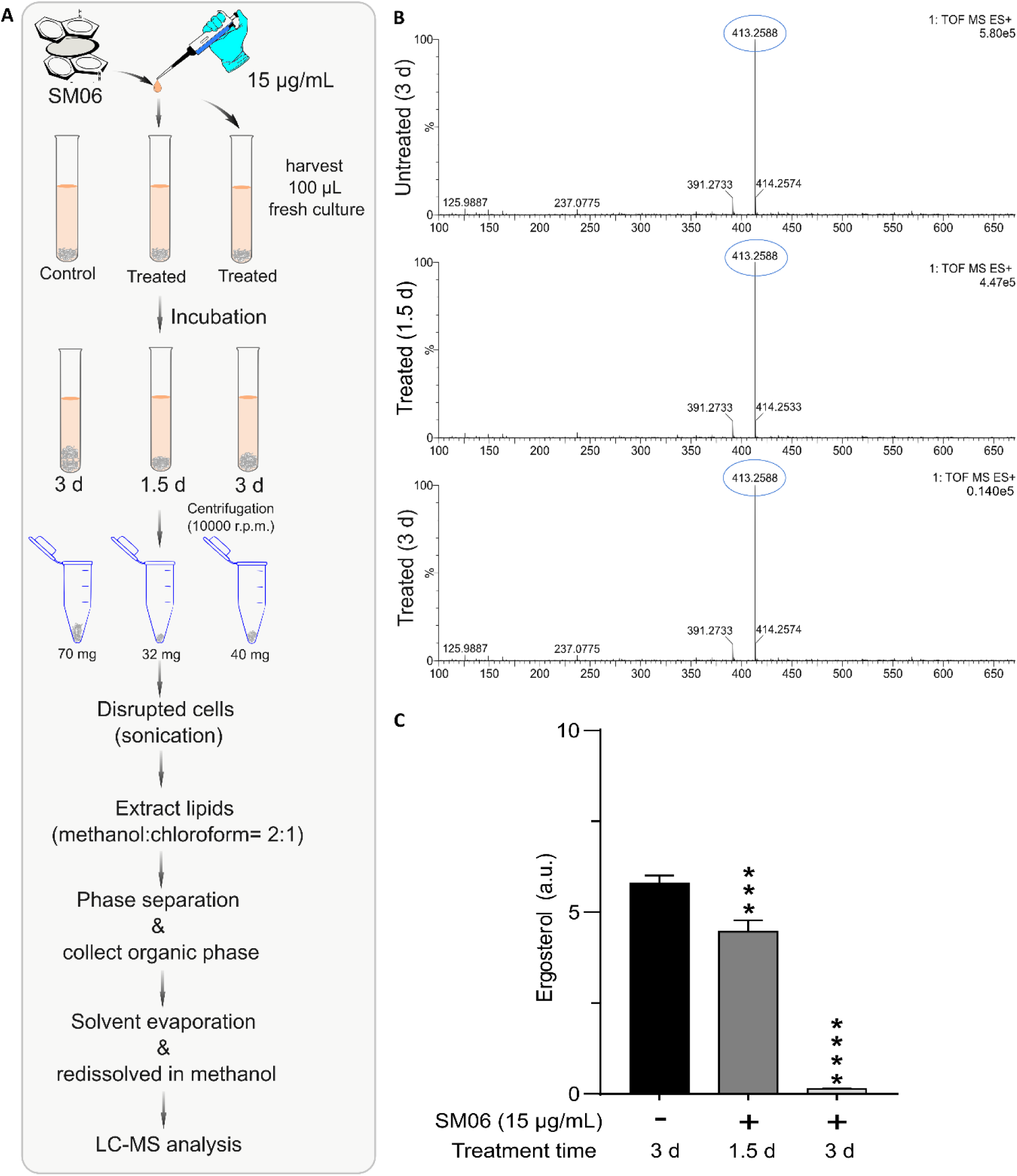
Time-dependent reduction of ergosterol upon SM06 treatment. A: Workflow of ergosterol extraction and LC–MS analysis from control and treated cultures harvested at 1.5 and 3 days. B: LC–MS spectra showing the ergosterol ion (m/z 413.2588) in untreated and treated samples, indicating a marked decrease after treatment, especially at 3 days. C: Quantified ergosterol abundance demonstrating significant, time-dependent depletion of ergosterol in SM06 treated cells compared to control.

### SM06 cures plant from *Curvularia* pathogenesis and produced by RSE02 inside host plant

*Phytobacter* sp. RSE02 enters and colonized inside tomato plants infected with fungal pathogen *C. lunata*. Confocal microscopic analysis showed that the RFP-labelled RSE02 was primarily found along the perivascular and epidermal layers (Fig. S12). Active in-plant proliferation is indicated by these persistent fluorescence aggregates. In planta transmission and colonization was also established following time dependent counts of the labelled RSE02 from the various plant tissue [20]. Untreated plants showed significant wilting and chlorosis as the disease progressed (Fig. S13AII), while healthy controls showed no changes (Fig. S13AI). Remarkably, RSE02-treated pathogen-infected plants showed significant recovery, with recovered turgor, greenness, and nearly normal morphology (Fig. S13AIII).

The indole dimer metabolite at m/z 235.1252 [M+H]^+^ was identified by chemical profiling of the tissue extracts from RSE02-treated, infected plants (Fig. S13B). This characteristic matched the SM06 reference standard in terms of retention time, precise mass, and fragmentation pattern (Fig. S13C). The lack of endogenous synthesis was indicated by the absence of such signal from the tissue extract of plant not infected with RSE02. These findings indicate that RSE02 colonization in plants ensures endogenous accumulation of SM06 indole dimer metabolite, which could essentially help to reduce the disease symptoms caused by fungal pathogens.

### SM06 controls brown leaf spot of tomato and rice

This study was designed to assess the protective efficacy of SM06 against *C. lunata*-induced brown leaf spot disease in tomato plants and fruits. The preventive and curative activities of SM06 were evaluated by applying foliar sprays of the compound at its minimum inhibitory concentration (MIC) to tomato leaves and fruits exhibiting freshly induced lesions (Fig. 7). Pathogen inoculation resulted in severe foliar wilting and chlorosis (Fig. 7AII), whereas both preventive and post-infection SM06 treatments markedly reduced symptom severity and restored plant vigor (Fig. 7AIII–IV). Microscopic observations revealed an intact tissue architecture in the control plants (Fig. 7BI) and dense hyphal colonization in the infected samples (Fig. 7BII). Curative application of SM06 substantially restricted hyphal spread (Fig. 7BIII), and preventive treatment completely inhibited pathogen ingress (Fig. 7BIV).

**Fig. 7.**
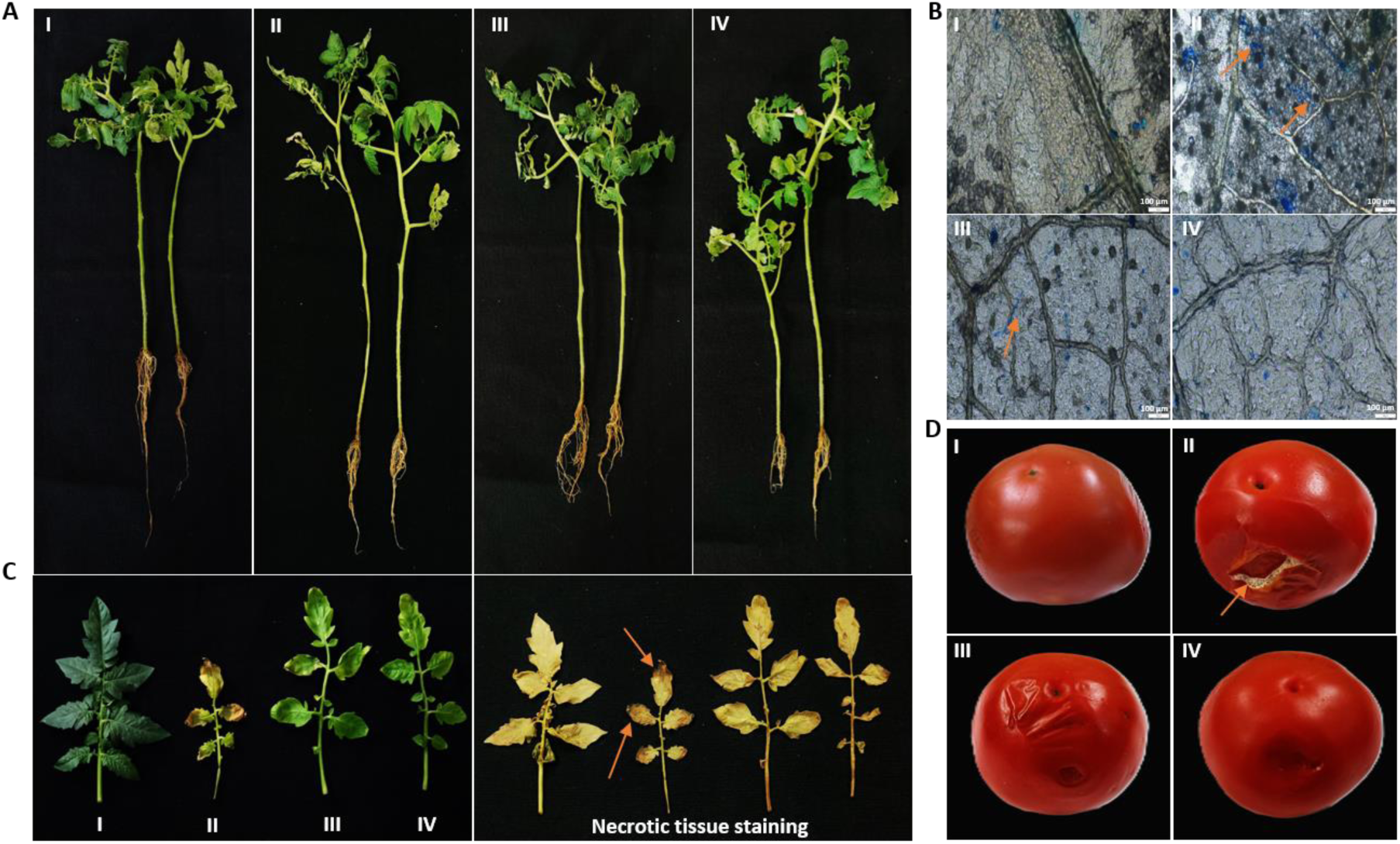
Antifungal activity of SM06 in *C. lunata* infected tomato plant. AI-IV: Images showing growth and infection parameter of experimental set of control, *C. lunata* infected plant, SM06 treated *C. lunata*-infected plant, and SM06 treated plant before *C. lunata* infection. BI-IV: Microscopic visualization of tomato leaf surface of the experimental sets A I-IV, respectively. The arrows (orange) indicate the colonization of *C. lunata* mycelium in the intercellular spaces on leaf (AI-IV). CI-IV: Images of the leaves of the plants from experimental sets of AI-IV; and DAB staining of tomato leaves to show the necrotic tissues. DI-IV: Effect of SM06 on control, *C. lunata* infected tomato, SM06 treated *C. lunata*-infected tomato, and SM06 treated tomato before *C. lunata* infection. All microscopic images are taken at 10X, and the bar shown in the images (AI-IV) is 100 μm.

Histochemical staining further supported these findings, as pathogen-infected leaves exhibited strong necrotic coloration (Fig. 7CI–II), whereas SM06-treated leaves showed minimal cell death (Fig. 7CIII–IV). Infection assays in fruits also confirmed reduced necrosis following SM06 application (Fig. 7DI–IV). Consistent with the results in tomato, SM06 also provide robust protection against brown leaf spot in rice, significantly limiting lesion formation and chlorosis (Fig. S14). Together, these results establish SM06 as a broad-spectrum antifungal agent with strong preventive and curative activities in multiple host species.

### RSE02 contains biosynthetic gene clusters for indole-like and other secondary metabolites

Genome mining revealed that *Phytobacter* sp. RSE02 encodes a diverse array of biosynthetic gene clusters (BGCs) linked to secondary metabolite formation (Fig. 8). Indole-related genes (*trpA, trpB,* and *tnaA*) were identified (Fig. 8A), along with multiple NRPS and PKS clusters predicted to synthesize aromatic polyketides and complex peptide metabolites.

**Fig. 8.**
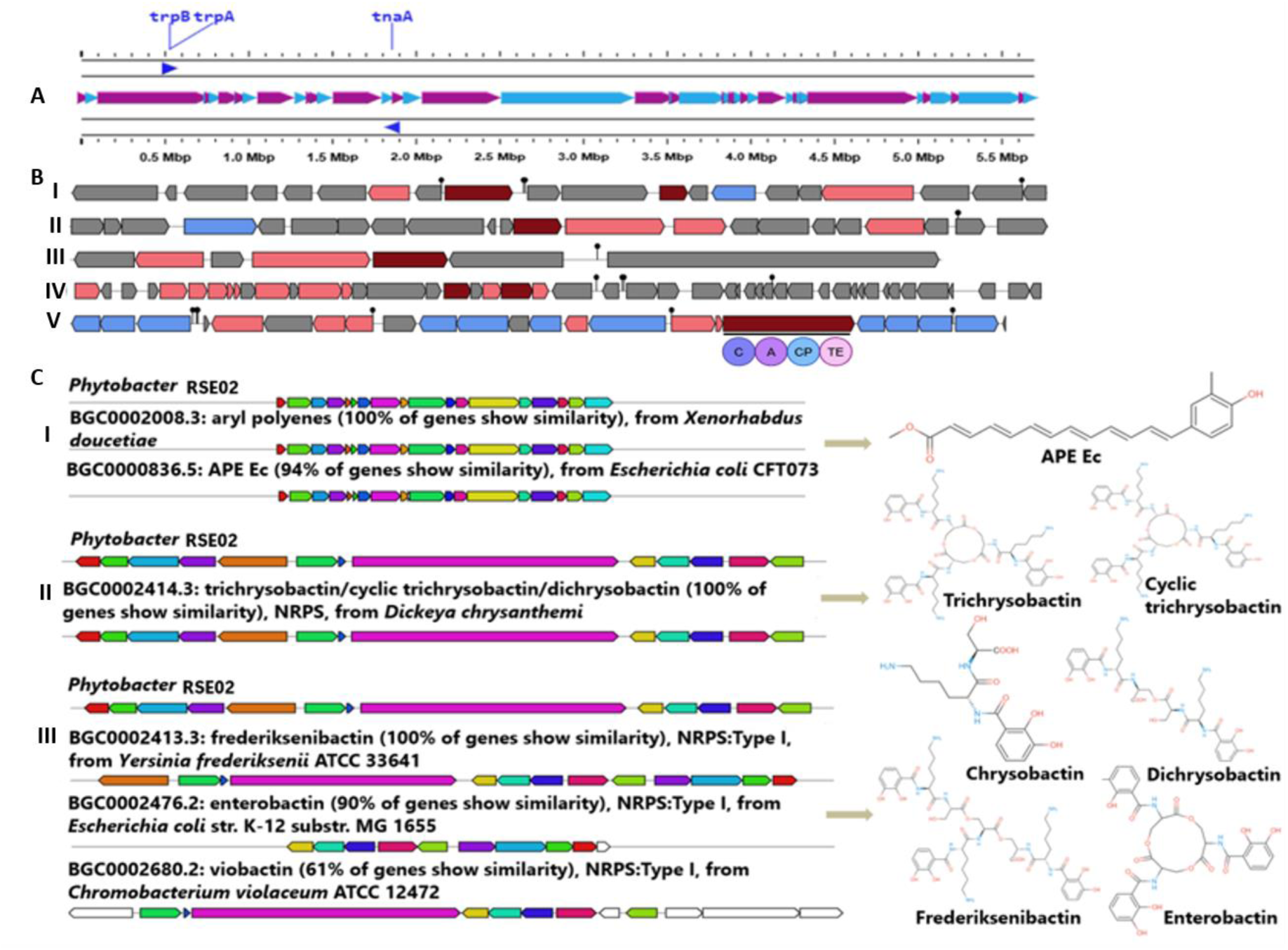
Biosynthetic gene clusters from RSE02 genome. A: Indole biosynthesis gene cluster (*trpA* & B, *tnaA*) in RSE02 genome. BI-V showing physical map of the biosynthetic gene clusters for azole-containing-RiPP, terpene-precursor, RiPP-like, arylpolyene and non-ribosomal peptide synthetase (NRPS), respectively. CI-III shows highly similar antimicrobial gene clusters of RSE02 compared with known clusters in the antiSMASH database. In the right panels the secondary metabolites are predicted to synthesized form the respective gene clusters.

AntiSMASH analysis showed presence of five biosynthetic gene clusters (BGCs) in the genome of *Phytobacter* sp. Strain RSE02. Clusters are found to be representing various secondary metabolites producing genes (Fig. 8BI-V) such as azole-containing-RiPP, terpene-precursor, RiPP-like, arylpolyene and non-ribosomal peptide synthetase (NRPS).

*Genome of Phytobacter* sp. strain RSE02 harbours BGC (Fig. 8CI) which showed 100% and 94% similarity with gene cluster producing aryl polyene and APE Ec (aryl polyene from *E. coli)* respectively. Additionally, the genome encodes siderophore biosynthetic gene cluster (Fig. 8CII, III) exhibiting 100% sequence identity to the NRPS-dependent cluster responsible for trichrysobactin, cyclic trichrysobactin, chrysobactin, and dichrysobactin biosynthesis whereas 90% similarity to the enterobactin gene cluster and 61% similarity to the viobactin gene cluster. This metabolic architecture indicates that RSE02 possesses a strong biosynthetic capacity for producing indole derivatives and additional bioactive compounds, with the diversity of its biosynthetic gene clusters suggesting a significant potential to synthesize a broad spectrum of metabolites consistent with its observed antifungal and plant beneficial phenotypes.

## DISCUSSION

The present study identified *Phytobacter* sp. RSE02 is a seed-associated ‘plant probiotic’ endophyte that produces a previously uncharacterized indole dimer antifungal metabolite, SM06, and establishes its broad-spectrum efficacy against diverse plant and clinical fungal pathogens. In contrast to our findings, a previous study demonstrated that di-indole metabolites derived from commensal microbiota thereby modulating host xenobiotic and metabolic pathways [48]. Our mechanistic, structural, and ecological analyses collectively indicate that SM06 acts primarily through competitive inhibition of fungal lanosterol 14-α-demethylase (ERG11) which cause perturbation of ergosterol balance and ultimately leading to impaired membrane integrity and fungal cell death.

### Mechanistic basis of SM06 antifungal action

Multiple lines of evidence indicate that SM06 targets the fungal cell membrane as its primary site of action. Enhanced propidium iodide (PI) uptake following SM06 treatment indicated loss of membrane integrity, consistent with necrotic cell death and in line with the established behavior of membrane-disruptive antifungals [49]. Scanning electron microscopy further revealed severe morphological aberrations, including hyphal swelling, septal distortion, and conidial surface collapse, consistent with catastrophic membrane and cell wall destabilization [50]. The thermal stability of SM06 across a temperature range (30°C-90°C) suggests that its antifungal activity is not easily abrogated, making it a promising candidate for various applications. Additionally, its broad-spectrum activity against diverse fungal pathogens underscores its potential as a versatile antifungal agent. Protein leakage assays revealed a time-dependent increase in extracellular protein content following SM06 treatment, indicating progressive disruption of the fungal cell membrane. The time-dependent leakage of intracellular proteins reinforces the conclusion that SM06 progressively compromises membrane permeability, a hallmark of lytic antifungal mechanisms [51]. From a detailed LC-MS analysis, it has also been found that there is significant reduction in fungal ergosterol levels (m/z 413.2588) upon SM06 treatment, strongly support the interference with sterol biosynthesis [52]. Ergosterol depletion is functionally congruent with the observed morphological abnormalities, given the central role of sterols in maintaining fungal membrane fluidity and structural order [53, 54]. Molecular docking further supports this mechanism, revealing that SM06 exhibits competitive binding within the catalytic pocket of ERG11, partially overlapping with the binding surfaces of both lanosterol and fluconazole. This mode of inhibition parallels the action of azoles but involves distinct structural interactions that may circumvent conventional azole resistance pathways, including ERG11 mutations and efflux activation [55, 56]. The partial synergism observed in combination assays with nystatin (FICI = 0.60) aligns with their complementary biochemical activities, SM06 disrupting sterol biosynthesis and Nystatin binding to ergosterol to further destabilize the membrane structure [57]. Collectively, these results establish that SM06 disrupts fungal cell integrity through dual mechanisms: direct membrane permeabilization and impairment of ergosterol biosynthesis.

### Structure–function comparisons with known antifungal indoles

The superior performance of SM06 in comparison to monomeric indoles can be rationalized by structural considerations. Prior studies have highlighted that dimeric or conjugated indole architectures exhibit enhanced biological potency due to increased hydrophobicity and improved target engagement [58, 59]. Similar patterns are reflected in synthetic 1,3,4-thiadiazole-indoles with strong activity against plant pathogens [60] and in dimeric indole-diterpenoids derived from *Penicillium* species that display strong receptor interactions [61]. In contrast, monomeric indoles exhibit moderate antifungal effects [62], suggesting that dimerization confers superior bioactivity and metabolic stability to the resulting dimers. Molecular docking suggested that within the fungal ERG11 active site, the two indole rings preferentially adopt a π–π–stacked configuration, further stabilised by hydrogen-bonding interactions between the N–H group of the dimer and key amino acid residues. This aligns with our ADMET profiling, which indicated improved lipophilicity, plasma protein binding, and extended half-life of SM06 relative to that of normal indole. The enhanced affinity of SM06 for ERG11 compared to fluconazole further underscores the functional advantage imparted by this dimeric architecture [63, 64].

### Ecological relevance and implications for plant protection

Endophytic bacteria that synthesize antifungal metabolites represent an important ecological strategy for suppressing plant diseases [65, 66]. Consistent with this paradigm, *Phytobacter* sp. RSE02 not only inhibited fungal pathogens in vitro but also conferred robust protection to tomato and rice plants against *C. lunata* infections. Preventive application of SM06 completely blocked fungal colonization, whereas curative application significantly limited lesion expansion and reduced the spread of hyphae. These findings parallel previous reports demonstrating the antifungal activities of small molecules against *Curvularia* sp. [67, 68] and expand the repertoire of endophyte-derived metabolites with agricultural potential.

The ability of RSE02 to colonize in plant tissues, as confirmed by confocal microscopy, further suggests that the endogenous production of SM06 within host tissues contributes to disease mitigation. The detection of SM06-like signatures (m/z 235.1252) in infected plants treated with RSE02 supports in planta biosynthesis or accumulation of this compound. This phenomenon is consistent with the established ecological roles of endophytes, which routinely synthesize protective metabolites under pathogen-challenge conditions [69, 70]. Beyond disease suppression, SM06 exhibited preservative benefits in tomato fruits, reducing infection-induced necrosis and oxidative damage. These effects align with the known natural preservative compounds in tomatoes (e.g., carotenoids and phenolic acids) [71], and support the utility of SM06 as a post-harvest protective agent [72].

### Genomic context and metabolic potential of RSE02

Genome mining of RSE02 revealed multiple biosynthetic gene clusters (BGCs), including tryptophan-derived indole pathways (*trpA*, *trpB*, and *tnaA*) and NRPS/PKS clusters (Fig. 8). This metabolic repertoire is characteristic of plant-associated Enterobacterales and is often linked to antifungal activity, competition, and beneficial host traits [73, 74]. The presence of indole biosynthetic genes supports the endogenous synthesis of indole dimer metabolites, such as SM06, while NRPS/PKS clusters suggest additional cryptic metabolites that may synergize with SM06 or exert independent protective effects.

## CONCLUSION

Overall, our results position *Phytobacter* sp. RSE02 and its indole dimer metabolite, SM06, are potent fungal antagonists with mechanistic, structural, and ecological features well-suited for agricultural and biotechnological applications. SM06 couples membrane-disruptive activity with targeted inhibition of ergosterol biosynthesis, conferring broad antifungal efficacy, including strong activity against *Curvularia* pathogens in multiple crop systems. The capacity of RSE02 to colonize host tissues and synthesize SM06 in planta highlights a promising route for eco-friendly, endophyte-based disease management, which aligns with global efforts to reduce dependence on synthetic fungicides.

## Ethical approval

This study was approved by the animal ethics committee (No. IAEC-04/BIOCHEM/CU/02-2022/03-SH) of Department of Biochemistry, University of Calcutta, INDIA.

## Acknowledgements

The authors sincerely acknowledge the experimental help provided by Priyabrata Mondal during antifungal studies. We also thank Kalipada Manna and Anirban Dolai, IIT Dhanbad for helping with the LC-MS data analysis. We are also thankful to DBT-Builder facility for Confocal Microscopy study and to the Department of Microbiology, University of Calcutta for providing research facilities.

## Author contributions

**Santosh Kumar Jana**: Writing – original draft, Writing – review & editing, Visualization, Methodology. **Supriya Bhunia**: Writing – review & editing, Visualization, Methodology. **Dipanjan Ghosh**: Writing – review & editing, Methodology. **Debashmita Guha**: Writing – review & editing, Methodology. **Shrodha Mondal**: Writing – review & editing, Methodology. **Himadri Sekhar Sarkar**: Methodology, Writing – review & editing. **Samudra Gupta**: Visualization, Methodology. **Subhas Samanta**: Writing – review & editing. **Prithidipa Sahoo**: Methodology, Writing – review & editing. **Sukhendu Mandal**: Conceptualization, Methodology, Writing – review & editing.

## Data availability

All relevant data are within the paper.

## Supporting Information

The data that support the findings of this study are available in the Supporting Information (SI) of this article.

## REFERENCE

1. Foley JA, Ramankutty N, Brauman KA, Cassidy ES, Gerber JS, Johnston M, Mueller ND, O’Connell C, Ray DK, West PC, Balzer C. 2011. Solutions for a cultivated planet. Nature 478:337–342. 10.1038/nature10452.

2. Fisher MC, Henk DA, Briggs CJ, Brownstein JS, Madoff LC, McCraw SL, Gurr SJ. 2012. Emerging fungal threats to animal, plant and ecosystem health. Nature 484:186–194. 10.1038/nature10947.

3. Saxena S, Dufossé L, Deshmukh SK, Chhipa H, Gupta MK. 2024. Endophytic fungi: a treasure trove of antifungal metabolites. Microorganisms 12:1903. 10.3390/microorganisms12091903.

4. Avenot HF, Morgan DP, Quattrini J, Michailides TJ. 2020. Resistance to thiophanate-methyl in *Botrytis cinerea* isolates from Californian vineyards and pistachio and pomegranate orchards. Plant Dis 104:1069–1075. 10.1094/PDIS-02-19-0353-RE.

5. Pathak VM, Verma VK, Rawat BS, Kaur B, Babu N, Sharma A, Dewali S, Yadav M, Kumari R, Singh S, Mohapatra A. 2022. Current status of pesticide effects on environment, human health, and its eco-friendly management as bioremediation: a comprehensive review. Front Microbiol 13:962619. 10.3389/fmicb.2022.962619.

6. Popp J, Pető K, Nagy J. 2013. Pesticide productivity and food security: a review. Agron Sustain Dev 33:243–255. 10.1007/s13593-012-0105-x.

7. Yin Y, Miao J, Shao W, Liu X, Zhao Y, Ma Z. 2023. Fungicide resistance: progress in understanding mechanism, monitoring, and management. Phytopathology 113:707–718. 10.1094/PHYTO-10-22-0370-KD.

8. Glick BR. 2012. Plant growth-promoting bacteria: mechanisms and applications. Scientifica 2012:963401. 10.6064/2012/963401.

9. Compant S, Duffy B, Nowak J, Clément C, Barka EA. 2005. Use of plant growth-promoting bacteria for biocontrol of plant diseases: principles, mechanisms of action, and future prospects. Appl Environ Microbiol 71:4951–4959. 10.1128/AEM.71.9.4951-4959.2005.

10. Madhaiyan M, Poonguzhali S, Lee JS, Saravanan VS, Lee KC, Santhanakrishnan P. 2010. *Enterobacter arachidis* sp. nov., a plant-growth-promoting diazotrophic bacterium isolated from rhizosphere soil of groundnut. Int J Syst Evol Microbiol 60:1559–1564. 10.1099/ijs.0.013664-0.

11. Pretty J, Bharucha ZP. 2015. Integrated pest management for sustainable intensification of agriculture in Asia and Africa. Insects 6:152–182. 10.3390/insects6010152.

12. Lugtenberg B, Kamilova F. 2009. Plant-growth-promoting rhizobacteria. Annu Rev Microbiol 63:541–556. 10.1146/annurev.micro.62.081307.162918.

13. Gond SK, Bergen MS, Torres MS, White JF Jr. 2015. Endophytic *Bacillus* spp. produce antifungal lipopeptides and induce host defence gene expression in maize. Microbiol Res 172:79–87. 10.1016/j.micres.2014.11.004.

14. Pandey SS, Jain R, Bhardwaj P, Thakur A, Kumari M, Bhushan S, Kumar S. 2022. Plant probiotics endophytes pivotal to plant health. Microbiol Res 263:127148. 10.1016/j.micres.2022.127148.

15. Xu K, Li XQ, Zhao DL, Zhang P. 2021. Antifungal secondary metabolites produced by fungal endophytes: chemical diversity and potential use in the development of biopesticides. Front Microbiol 12:689527. 10.3389/fmicb.2021.689527.

16. Debnath S, Addya S. 2014. Structural basis for heterogeneous phenotype of ERG11-dependent azole resistance in *Candida albicans* clinical isolates. Springerplus 3:660. 10.1186/2193-1801-3-660.

17. Prakash SM, Nazeer Y, Jayanthi S, Kabir MA. 2020. Computational insights into fluconazole resistance by suspected mutations in lanosterol 14α-demethylase (Erg11p) of *Candida albicans*. Mol Biol Res Commun 9:155. 10.22099/mbrc.2020.36298.1476.

18. Choi JY, Podust LM, Roush WR. 2014. Drug strategies targeting CYP51 in neglected tropical diseases. Chem Rev 114:11242–11271. 10.1021/cr5003134.

19. Sharma A, Kaushik N, Sharma A, Marzouk T, Djébali N. 2022. Exploring the potential of endophytes and their metabolites for biocontrol activity. 3 Biotech 12:277. 10.1007/s13205-022-03321-0.

20. Jana SK, Islam MM, Hore S, Mandal S. 2023. Rice seed endophytes transmit into the plant seedling, promote plant growth, and inhibit fungal phytopathogens. Plant Growth Regul 99:373–388. 10.1007/s10725-022-00914-w.

21. Maiti PK, Das S, Sahoo P, Mandal S. 2020. *Streptomyces* sp. SM01 isolated from Indian soil produces a novel antibiotic picolinamycin effective against multidrug-resistant bacterial strains. Sci Rep 10:10092. 10.1038/s41598-020-66984-w.

22. Islam MM, Saha S, Sahoo P, Mandal S. 2024. Endophytic *Streptomyces* sp. MSARE05 isolated from roots of peanut produces a novel antimicrobial compound. J Appl Microbiol 135:lxae051. 10.1093/jambio/lxae051.

23. Ruhil S, Balhara M, Dhankhar S, Kumar M, Kumar V, Chhillar AK. 2013. Advancement in infection control of opportunistic pathogens (*Aspergillus* spp.): adjunctive agents. Curr Pharm Biotechnol 14:226–232. 10.2174/138920113805219359.

24. Sharma M, Manhas RK. 2020. Purification and characterization of salvianolic acid B from *Streptomyces* sp. M4 possessing antifungal activity against fungal phytopathogens. Microbiol Res 237:126478. 10.1016/j.micres.2020.126478.

25. Bradley RL Jr. 2010. Moisture and total solids analysis. In Nielsen SS (ed), Food analysis, p 85–104. Springer, Boston, MA. 10.1007/978-1-4419-1478-1_6.

26. Ruhil S, Kumar V, Balhara M, Malik M, Dhankhar S, Kumar M, Chhillar AK. 2014. In vitro evaluation of combinations of polyenes with EDTA against *Aspergillus* spp. by different methods (FICI and CI model). J Appl Microbiol 117:643–653. 10.1111/jam.12576.

27. Banerjee N, Sengupta S, Roy A, Ghosh P, Das K, Das S. 2011. Functional alteration of a dimeric insecticidal lectin to a monomeric antifungal protein correlated to its oligomeric status. PLoS One 6:e18593. 10.1371/journal.pone.0018593.

28. Mani-López E, Cortés-Zavaleta O, López-Malo A. 2021. Methods used to determine the target site or mechanism of action of essential oils and their components against fungi. SN Appl Sci 3:44. 10.1007/s42452-020-04102-1.

29. Bradford MM. 1976. A rapid and sensitive method for the quantitation of microgram quantities of protein utilizing the principle of protein-dye binding. Anal Biochem 72:248–254. 10.1016/0003-2697(76)90527-3.

30. Sardana K, Gupta A, Sadhasivam S, Gautam RK, Khurana A, Saini S, Gupta S, Ghosh S. 2021. Checkerboard analysis to evaluate synergistic combinations of existing antifungal drugs and propylene glycol monocaprylate in isolates from recalcitrant tinea corporis and cruris harboring squalene epoxidase gene mutation. Antimicrob Agents Chemother 65:e00321–21. 10.1128/AAC.00321-21.

31. Rea V, Bell I, Ball T, Van Raay T. 2022. Gut-derived metabolites influence neurodevelopmental gene expression and Wnt signaling events in a germ-free zebrafish model. Microbiome 10:132. 10.1186/s40168-022-01302-2.

32. Mukherjee S, Das S, Das S, Gupta S, Hui SP, Sengupta A, Ghosh A. 2025. Pyruvate plus uridine augments mitochondrial respiration and prevents cardiac hypertrophy in zebrafish and H9c2 cells. J Cell Sci 138:jcs263653. 10.1242/jcs.263653.

33. Mosmann T. 1983. Rapid colorimetric assay for cellular growth and survival: application to proliferation and cytotoxicity assays. J Immunol Methods 65:55–63. 10.1016/0022-1759(83)90303-4.

34. Liu Y, Grimm M, Dai WT, Hou MC, Xiao ZX, Cao Y. 2020. CB-Dock: a web server for cavity detection-guided protein ligand blind docking. Acta Pharmacol Sin 41:138–144. 10.1038/s41401-019-0228-6.

35. Hanwell MD, Curtis DE, Lonie DC, Vandermeersch T, Zurek E, Hutchison GR. 2012. Avogadro: an advanced semantic chemical editor, visualization, and analysis platform. J Cheminform 4:17. 10.1186/1758-2946-4-17.

36. El-Hachem N, Haibe-Kains B, Khalil A, Kobeissy FH, Nemer G. 2017. AutoDock and AutoDockTools for protein-ligand docking: beta-site amyloid precursor protein cleaving enzyme 1 (BACE1) as a case study. In Kobeissy FH (ed), Neuroproteomics: methods and protocols, p 391–403. Springer, New York, NY. 10.1007/978-1-4939-6952-4_20.

37. Yuan S, Chan HS, Hu Z. 2017. Using PyMOL as a platform for computational drug design. WIREs Comput Mol Sci 7:e1298. 10.1002/wcms.1298.

38. Swanson K, Walther P, Leitz J, Mukherjee S, Wu JC, Shivnaraine RV, Zou J. 2024. ADMET-AI: a machine learning ADMET platform for evaluation of large-scale chemical libraries. Bioinformatics 40:btae416. 10.1093/bioinformatics/btae416.

39. Wilkes TI. 2023. Ergosterol extraction: a comparison of methodologies. Access Microbiol 5:000490. 10.1099/acmi.0.000490.v4.

40. Jana SK, Bhattacharya R, Mukherjee S, Gupta S, Hui SP, Chattopadhyay A, Biswas SR, Mandal S. 2025. *Phytobacter* sp. RSE02 is a rice seed endophytic plant probiotic bacterium with human probiotic features and cholesterol-lowering ability. Sci Rep 15:27865. 10.1038/s41598-025-11212-6.

41. Revelou PK, Kokotou MG, Constantinou-Kokotou V. 2020. Determination of indole-type phytonutrients in cruciferous vegetables. Nat Prod Res 34:2554–2557. 10.1080/14786419.2018.1543680.

42. Wang Z, Xia Y, Lin S, Wang Y, Guo B, Song X, Ding S, Zheng L, Feng R, Chen S, Bao Y. 2018. Osa-miR164a targets OsNAC60 and negatively regulates rice immunity against the blast fungus *Magnaporthe oryzae*. Plant J 95:584–597. 10.1111/tpj.13972.

43. Limtong S, Into P, Attarat P. 2020. Biocontrol of rice seedling rot disease caused by *Curvularia lunata* and *Helminthosporium oryzae* by epiphytic yeasts from plant leaves. Microorganisms 8:647. 10.3390/microorganisms8050647.

44. Namburi KR, Kora AJ, Chetukuri A, Kota VS. 2021. Biogenic silver nanoparticles as an antibacterial agent against bacterial leaf blight-causing rice phytopathogen *Xanthomonas oryzae* pv. *oryzae*. Bioprocess Biosyst Eng 44:1975–1988. 10.1007/s00449-021-02579-7.

45. Yadav U, Anand V, Kumar S, Verma I, Anshu A, Pandey IA, Kumar M, Behera SK, Srivastava S, Singh PC. 2024. *Bacillus subtilis* NBRI-W9 simultaneously activates SAR and ISR against *Fusarium chlamydosporum* NBRI-FOL7 to increase wilt resistance in tomato. J Appl Microbiol 135:lxae013. 10.1093/jambio/lxae013.

46. Roschzttardtz H, Conéjéro G, Curie C, Mari S. 2009. Identification of the endodermal vacuole as the iron storage compartment in the *Arabidopsis* embryo. Plant Physiol 151:1329–1338. 10.1104/pp.109.144444.

47. Maiti PK, Mandal S. 2022. Comprehensive genome analysis of *Lentzea* reveals a repertoire of polymer-degrading enzymes and bioactive compounds with clinical relevance. Sci Rep 12:8409. 10.1038/s41598-022-12427-7.

48. Liu J, Malekoltojari A, Asokakumar A, Chow V, Li L, Li H, Grimaldi M, Dang N, Campbell J, Barrett H, Sun J. 2024. Diindoles produced from commensal microbiota metabolites function as endogenous CAR/Nr1i3 ligands. Nat Commun 15:2563. 10.1038/s41467-024-46559-3.

49. Yu X, Yang S, Su P, Bi H, Li Y, Peng X, Sun X, Wang Q. 2025. 4-Propylphenol alters membrane integrity in fungi isolated from walnut anthracnose and brown spot. J Fungi 11:610. 10.3390/jof11090610.

50. Mercer DK, Stewart CS, Miller L, Robertson J, Duncan VM, O’Neil DA. 2019. Improved methods for assessing therapeutic potential of antifungal agents against dermatophytes and their application in the development of NP213, a novel onychomycosis therapy candidate. Antimicrob Agents Chemother 63:e02117–18. 10.1128/AAC.02117-18.

51. Struyfs C, Cammue BPA, Thevissen K. 2021. Membrane-interacting antifungal peptides. Front Cell Dev Biol 9:649875. 10.3389/fcell.2021.649875.

52. Ory L, Gentil E, Kumla D, Kijjoa A, Nazih EH, Roullier C. 2020. Detection of ergosterol using liquid chromatography electrospray ionization mass spectrometry: investigation of unusual in-source reactions. Rapid Commun Mass Spectrom 34:e8780. 10.1002/rcm.8780.

53. Ostrosky-Zeichner L, Casadevall A, Galgiani JN, Odds FC, Rex JH. 2010. An insight into the antifungal pipeline: selected new molecules and beyond. Nat Rev Drug Discov 9:719–727. 10.1038/nrd3074.

54. Nishimoto AT, Sharma C, Rogers PD. 2020. Molecular and genetic basis of azole antifungal resistance in the opportunistic pathogenic fungus *Candida albicans*. J Antimicrob Chemother 75:257–270. 10.1093/jac/dkz400.

55. Cowen LE, Sanglard D, Howard SJ, Rogers PD, Perlin DS. 2015. Mechanisms of antifungal drug resistance. Cold Spring Harb Perspect Med 5:a019752. 10.1101/cshperspect.a019752.

56. Sanglard D. 2016. Emerging threats in antifungal-resistant fungal pathogens. Front Med 3:11. 10.3389/fmed.2016.00011.

57. Cui J, Ren B, Tong Y, Dai H, Zhang L. 2015. Synergistic combinations of antifungals and anti-virulence agents to fight against *Candida albicans*. Virulence 6:362–371. 10.1080/21505594.2015.1039885.

58. Zou H, Wang Y, Pu H, Fu H, Cheng Z, Tian F, Gu L, Xue W. 2025. Discovery of novel indole derivatives containing imidazolidinone as potential antifungal agents with mechanistic studies. J Agric Food Chem 73:27363–27371. 10.1021/acs.jafc.5c09130.

59. Niu J, Qi J, Wang P, Liu C, Gao JM. 2023. Chemical structures and biological activities of indole diterpenoids. Nat Prod Bioprospect 13:3. 10.1007/s13659-022-00368-7.

60. He B, Hu Y, Xing L, Qing Y, Meng K, Zeng W, Sun Z, Wang Z, Xue W. 2024. Antifungal activity of novel indole derivatives containing 1,3,4-thiadiazole. J Agric Food Chem 72:10227–10235. 10.1021/acs.jafc.3c09303.

61. Du HF, Li L, Zhang YH, Wang X, Zhou CY, Zhu HJ, Pittman CU, Shou JW, Cao F. 2025. The first dimeric indole-diterpenoids from a marine-derived *Penicillium* sp. fungus and their potential for anti-obesity drugs. Mar Life Sci Technol 7:120–131. 10.1007/s42995-024-00253-x.

62. Ryu CK, Lee JY, Park RE, Ma MY, Nho JH. 2007. Synthesis and antifungal activity of 1H-indole-4,7-diones. Bioorg Med Chem Lett 17:127–131. 10.1016/j.bmcl.2006.09.076.

63. Song JL, Harry JB, Eastman RT, Oliver BG, White TC. 2004. The *Candida albicans* lanosterol 14α-demethylase (ERG11) gene promoter is maximally induced after prolonged growth with antifungal drugs. Antimicrob Agents Chemother 48:1136–1144. 10.1128/AAC.48.4.1136-1144.2004.

64. Flowers SA, Colón B, Whaley SG, Schuler MA, Rogers PD. 2015. Contribution of clinically derived mutations in ERG11 to azole resistance in *Candida albicans*. Antimicrob Agents Chemother 59:450–460. 10.1128/AAC.03470-14.

65. Pieterse CMJ, Zamioudis C, Berendsen RL, Weller DM, Van Wees SCM, Bakker PAHM. 2014. Induced systemic resistance by beneficial microbes. Annu Rev Phytopathol 52:347–375. 10.1146/annurev-phyto-082712-102340.

66. Hardoim PR, Van Overbeek LS, Berg G, Pirttilä AM, Compant S, Campisano A, Döring M, Sessitsch A. 2015. The hidden world within plants: ecological and evolutionary considerations for defining functioning of microbial endophytes. Microbiol Mol Biol Rev 79:293–320. 10.1128/MMBR.00050-14.

67. Chang X, Wang Y, Zain A, Yu H, Huang W. 2024. Antifungal activity of difenoconazole-loaded microcapsules against *Curvularia lunata*. J Fungi 10:519. 10.3390/jof10080519.

68. Hajji-Hedfi L, Rhouma A, Wannassi T, Utkina AO, Rebouh NY. 2025. Biocontrol assessment of *Trichoderma* species on tomato crops infested by *Curvularia spicifera*: towards sustainable farming systems. Front Sustain Food Syst 9:1627903. 10.3389/fsufs.2025.1627903.

69. Rodriguez RJ, White JF Jr, Arnold AE, Redman AR. 2009. Fungal endophytes: diversity and functional roles. New Phytol 182:314–330. 10.1111/j.1469-8137.2009.02773.x.

70. Trivedi P, Leach JE, Tringe SG, Sa T, Singh BK. 2020. Plant–microbiome interactions: from community assembly to plant health. Nat Rev Microbiol 18:607–621. 10.1038/s41579-020-0412-1.

71. Chaudhary P, Sharma A, Singh B, Nagpal AK. 2018. Bioactivities of phytochemicals present in tomato. J Food Sci Technol 55:2833–2849. 10.1007/s13197-018-3221-z.

72. Zaman W, Amin A, Khalil AA, Akhtar MS, Ali S. 2025. Plant–microbe interactions for improving postharvest shelf life and quality of fresh produce through protective mechanisms. Horticulturae 11:732. 10.3390/horticulturae11070732.

73. Medema MH, Fischbach MA. 2015. Computational approaches to natural product discovery. Nat Chem Biol 11:639–648. 10.1038/nchembio.1884.

74. Carrión VJ, Perez-Jaramillo J, Cordovez V, Tracanna V, De Hollander M, Ruiz-Buck D, Mendes LW, van Ijcken WFJ, Gomez-Exposito R, Elsayed SS, Mohanraju P. 2019. Pathogen-induced activation of disease-suppressive functions in the endophytic root microbiome. Science 366:606–612. 10.1126/science.aaw9285.

